# Single-cell RNA-sequencing reveals profibrotic roles of distinct epithelial and mesenchymal lineages in pulmonary fibrosis

**DOI:** 10.1101/753806

**Authors:** Arun C. Habermann, Austin J. Gutierrez, Linh T. Bui, Stephanie L. Yahn, Nichelle I. Winters, Carla L. Calvi, Lance Peter, Mei-I Chung, Chase J. Taylor, Christopher Jetter, Latha Raju, Jamie Roberson, Guixiao Ding, Lori Wood, Jennifer MS Sucre, Bradley W. Richmond, Ana P. Serezani, Wyatt J. McDonnell, Simon B. Mallal, Matthew J. Bacchetta, James E. Loyd, Ciara M. Shaver, Lorraine B. Ware, Ross Bremner, Rajat Walia, Timothy S. Blackwell, Nicholas E. Banovich, Jonathan A. Kropski

## Abstract

Pulmonary fibrosis is a form of chronic lung disease characterized by pathologic epithelial remodeling and accumulation of extracellular matrix. In order to comprehensively define the cell types, mechanisms and mediators driving fibrotic remodeling in lungs with pulmonary fibrosis, we performed single-cell RNA-sequencing of single-cell suspensions from 10 non-fibrotic control and 20 PF lungs. Analysis of 114,396 cells identified 31 distinct cell types. We report a remarkable shift in epithelial cell phenotypes occurs in the peripheral lung in PF, and identify several previously unrecognized epithelial cell phenotypes including a *KRT5*^−^/*KRT17*^+^, pathologic ECM-producing epithelial cell population that was highly enriched in PF lungs. Multiple fibroblast subtypes were observed to contribute to ECM expansion in a spatially-discrete manner. Together these data provide high-resolution insights into the complexity and plasticity of the distal lung epithelium in human disease, and indicate a diversity of epithelial and mesenchymal cells contribute to pathologic lung fibrosis.

**One Sentence Summary:** Single-cell RNA-sequencing provides new insights into pathologic epithelial and mesenchymal remodeling in the human lung.

## Main Text

Pulmonary fibrosis (PF) is a form of chronic lung disease characterized by progressive accumulation of extracellular matrix (ECM) in the peripheral lung accompanied by destruction of functional alveolar gas-exchange units. The most severe form of PF, idiopathic pulmonary fibrosis (IPF), is a relentlessly progressive disorder of unknown cause that leads to respiratory failure and death or lung transplantation within 3-5 years of diagnosis. Spatially heterogeneous expansion of interstitial collagen and extracellular matrix (i.e. “temporal heterogeneity”) is a defining feature of the histopathology of IPF (*1*). Epithelial-mesenchymal interactions have been proposed as playing a central role driving pathologic epithelial remodeling and ECM expansion. While there have been advances in identifying factors that regulate fibrosis in experimental models, to date there has been limited progress toward developing an integrated understanding of the central mechanisms driving pathologic epithelial remodeling and ECM expansion in the human lung. Using single-cell RNA sequencing (scRNA-seq), we identified multiple distinct epithelial and mesenchymal lineages contributing to ECM expansion in pulmonary fibrosis lungs. In addition to myofibroblasts, we identified a unique *HAS1^hi^*, ECM-producing population which is markedly enriched in lungs from patients with IPF and localizes to peripheral and subpleural regions, and reported a previously undescribed *KRT5*^−^/*KRT17*^+^ epithelial cell population expressing collagen and other ECM components that is conserved across a subset of histopathologic patterns of PF. Unbiased analyses revealed a spectrum of pathologic epithelial cell programs in the distal lung, suggesting an unexpected degree of plasticity. Together these high-resolution transcriptomic data and the identification of multiple novel pathologic cell types provide remarkable insights into the plasticity of the human lung and the fundamental mechanisms driving disease pathology in PF.

In order to determine cellular populations and mediators shared across different forms of pulmonary fibrosis, we generated single-cell suspensions from peripheral lung tissue removed at the time of lung transplant surgery from patients with IPF (n = 12), chronic hypersensitivity pneumonitis (cHP; n = 3), nonspecific interstitial pneumonia (NSIP; n = 2), sarcoidosis (n = 2), unclassifiable interstitial lung disease (ILD; n = 1), as well as nonfibrotic controls (declined donors; n = 10) (Table S1) and performed scRNA-seq using the 10X Genomics Chromium platform (see Methods; Fig. 1A). Jointly analyzing data from all samples we recovered 114,396 cells and performed unsupervised clustering using the Seurat (*2, 3*) package in R, followed by manual annotation, which defined 31 distinct cell types in the lung (see Methods; Fig. 1B, C, Table S2-S3, Figs. S1-3). Comparing cells originating from PF lungs to controls, in each cell type hundreds genes were differentially expressed(see Methods; Fig. 1D, Table S4). We recapitulated known expression differences linked to PF (Fig. 1E), demonstrated novel transcriptional variation, and identified patterns of shared and cell type specific expression changes (Fig. S3). We identified multiple cell types expressing genes encoding for ECM components increased in IPF lungs (*4*). As expected, these ECM genes were largely expressed in fibroblast sub-populations, however expression of a subset of these genes was identified in smooth muscle cells, endothelial cells, AT1 and AT2 cells (Fig. 1F, Fig. S4). This analysis also identified a previously undescribed population of *KRT5*^−^/*KRT17*^+^, collagen-producing epithelial cells (Fig. 1F).

**Figure 1.**
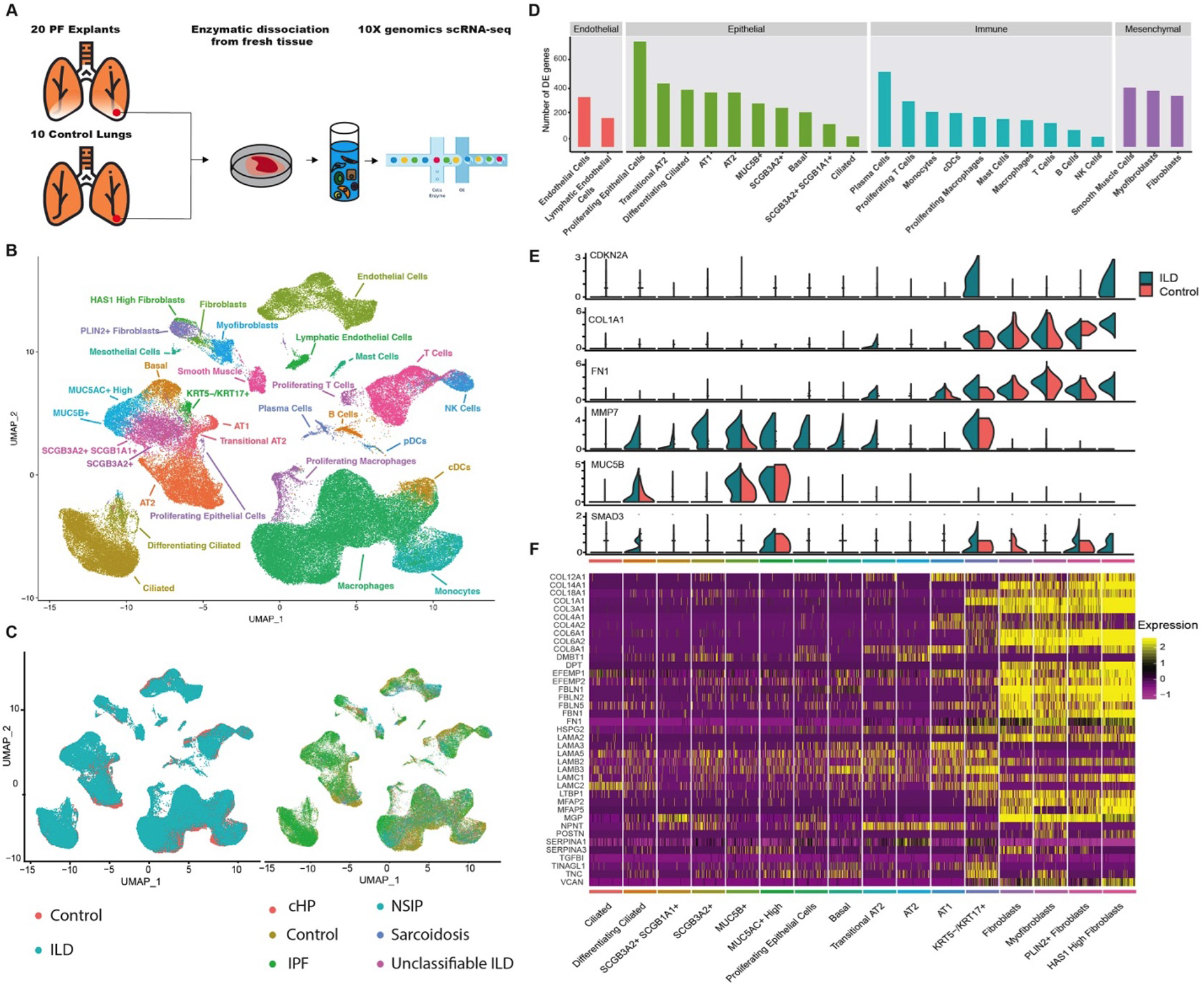
Single cell landscape of PF and control lungs. **A**) Schematic of workflow for scRNA-seq using the 10X Chromium platform. UMAP embedding of jointly analyzed single-cell transcriptomes from 114,396 cells from 20 PF and 10 control lungs annotated by **B**) cell type, **C**) disease status. **D**) Number of differentially expressed genes in each cell type with >50 cells available in PF and control lungs. **E**) Cell type of origin and disease-state informed expression of selected biomarkers and putative mediators of pulmonary fibrosis. **F**) Heatmap depicting relative expression of ECM components previously shown to be increased in pulmonary fibrosis lungs.

Genetic evidence has suggested a central role of epithelial cells in mediating risk for IPF (*5–8*). Within epithelial cells (Fig. 2A), we observed large-scale changes in cell proportions between PF and control lungs (Fig. 2B-C), and identified multiple previously undescribed epithelial cell phenotypes. In general, we found cells arising from the airway epithelium (*KRT5*^+^ Basal cells, *FOXJ1*^+^ Ciliated cells, and secretory cells) to be expanded and cells from the alveolar epithelium (AT2 and AT1 cells) to be less frequently recovered from PF lungs. In addition to a putatively differentiating state characterized by expression of both secretory and ciliated programs, we identified four distinct secretory cell phenotypes (Fig. 2B). Two of these secretory cells expressed classical secretory lineage marker *SCGB1A1* as well as airways mucins (*MUC5AC* and/or *MUC5B*). Two other secretory populations were defined by expression of *SCGB1A1* and *SCGB3A2*, or *SCGB3A2* only (Fig. 2B, Fig. S5).

**Figure 2.**
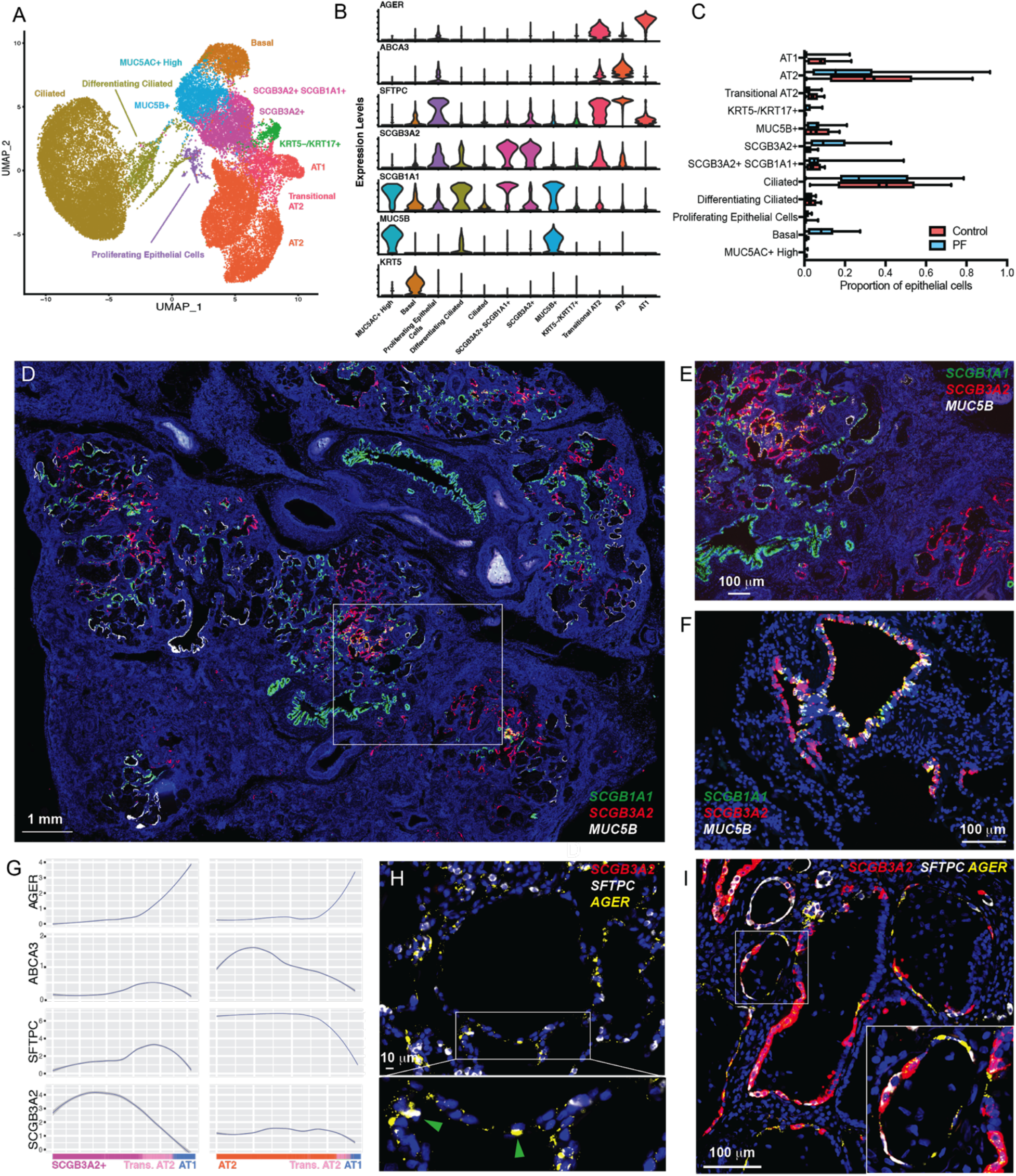
Epithelial cell identification in PF lungs. Cell types **A**) and canonical lineage markers **B**) identified in the peripheral lung. **C**) Quantification of cell types as a percent of all epithelial cells in PF vs. control lungs. Bxes indicate interquartile range and range. **D**) Low power stitched image of IPF lung labeling secretory lineages using Multiplexed RNA-ISH **E**) Higher magnification of box from D). **F**) RNA-ISH demonstrating secretory lineages in control lung. **G**) Smoothed expression of lineage markers along pseudotime trajectories from *SCGB3A2*^+^ and AT2 cells to AT1 from PF and control lungs (See figure S6). **H**) RNA-ISH demonstrating *AGER*^+^/*SFTPC*^+^ cells in control lungs **I**) and *AGER*^+^/*SFTPC*^+^/*SCGB3A2*^+^; in PF lungs.

To understand the spatial distribution of these secretory lineages within PF and control lungs, we performed RNA in-situ hybridization (ISH). In areas of cystic and fibrotic remodeling in PF lungs, we observed a striking pattern characterized by discrete regions of remodeled epithelium with a near exclusive population by a single secretory phenotype (Fig. 2D-E). In contrast, in control lungs, low-level *MUC5B* expression was observed in *SCGB1A1*^+^ cells in large airways; a subset of *SCGB1A1*^+^ cells coexpressed *SCGB3A2*. Occasional *SCGB3A2*^+^ cells lacking *SCGB1A1* or *MUC5B* expression were found in a subset of airways in control lungs (Fig. 2F). Consistent with our transcriptomic data, *SCGB3A2*^+^ epithelial cells were widespread in PF tissue sections, particularly in areas of cystic and fibrotic remodeling. Analysis of gene expression programs discriminating between the *SCGB3A2*^+^/*SCGB1A1*^+^ and *SCGB3A2*^+^ populations demonstrated the *SCGB3A2*^+^ cells expressed a subset of alveolar programs including *NKX2-1, HOPX, CAV1*, surfactant genes, and MHC-II (Fig. S5).

In both control and PF lungs, we identified a population of cells that express some features of both AT2 and AT1 cells (Fig. 2A, B), resembling an indeterminate population reported in previous work (*9*). Trajectory analysis performed using either mRNA processing or pseudotime (*10, 11*) (see Methods) suggested this Transitional AT2 cell population represents a state during the differentiation trajectory from AT2 to AT1 with increasing AT1 markers along the trajectory (Fig. 2G, S6). Surprisingly, among in both control and PF lungs, a proportion of Transitional AT2 cells express *SCGB3A2* (Fig. S7), and trajectory analyses demonstrated *SCGB3A2*^+^ cells upregulating AT1 and AT2 programs, leading us to hypothesize that *SCGB3A2*^+^ cells are capable of generating AT1 cells by differentiating to Transitional AT2 cells (Figure 2G). These results are consistent with a recent report suggesting both club cells and AT2 cells may generate AT1 cells via a common progenitor state in mice following experimental lung injury (*12*). To confirm these states exist *in-vivo*, we performed RNA-ISH using tissue sections from sequenced lungs to localize *SCGB3A2*, *SFTPC* (*AT2 marker*), and *AGER* (AT1 marker), and identified a putatively transitional state coexpressing *SFTPC* and *AGER* in both control (Fig. 2H) and fibrotic lungs (Fig. 2I); consistent with transcriptomic data, a subset of these *SFTPC^+^/AGER^+^* dual positive cells expressed low levels of *SCGB3A2* in PF samples and were rarely observed in control lungs (Fig. 2I). Together, these data indicate considerable plasticity contributing to the distal lung epithelial lineages, particularly in injured lungs, raising the possibility that multiple distal lineages converge upon a common transitional phenotype that can give rise to AT1 cells during injury-repair.

Surprisingly, we also identified a previously undescribed epithelial cell population that expressed *COL1A1* and other pathologic ECM components and was found nearly exclusively in PF lungs (Fig. 3A). Furthermore, these cells were identified across all histopathologic patterns of PF, though were infrequently recovered from lungs with NSIP pathology (Fig. S8). These *KRT5*^−^/*KRT17*^+^ cells expressed multiple collagens, other ECM components, and *CDH2* (N-cadherin), but lacked *ACTA2*, *PDGFRA*, *S100A4* or other canonical fibroblast markers (Fig. 1,3A). Although expressing some markers typical of basal cells, this *KRT5*^−^/*KRT17*^+^ population lacked *SOX2* expression, *KRT5* expression (Fig S9) and had high expression of genes typical of a distal epithelial program including *SOX9*, *NAPSA*, and *ITGB6* (Fig. 3A, Fig. S10, Table S5). Remarkably, this *KRT5*^−^/*KRT17*^+^ population highly expressed *MMP7*, the most well validated IPF biomarker in peripheral blood (*13*), was the predominant cell type expressing integrin *αVβ6*, previously implicated in transforming growthfactor β activation (*14*), and highly expressed *CDKN2A* (encoding for p16), suggesting cell-cycle arrest and/or a senescent phenotype (Fig. S10). Using RNA-ISH, we found *KRT5*^−^/*KRT17*^+^ cells intimately overlay foci of high collagen expression in the distal lung and coexpressed *COL1A1* (Fig. 3B-D); *KRT17*^+^ basal cells were observed in airways of control lungs but lacked co-expression of *COL1A1* (Fig. S9, S11). Furthermore, *KRT17*^+^/*COL1A1*^+^ cells were identified in surgical lung biopsy specimens from early stage IPF, and in transbronchial lung biopsy specimens from a cohort of asymptomatic individuals with a family history of IPF (*15*), indicating this unique cell type is present during early disease pathogenesis (Fig. 3E).

**Figure 3.**
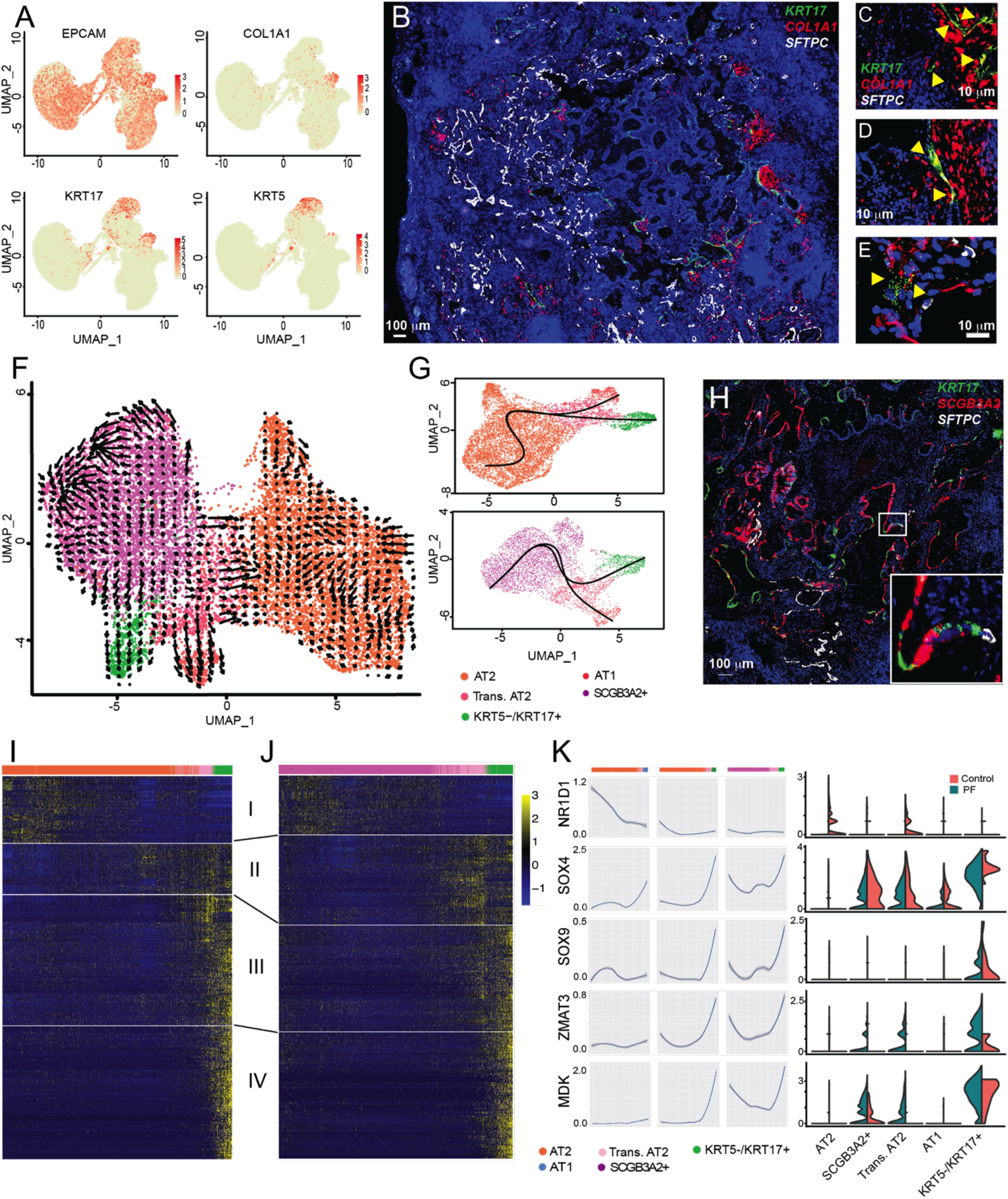
*KRT5*^−^/*KRT17*^+^ epithelial cells emerge in PF lungs. **A**) Identification of a *COL1A1*-expressing epithelial population characterized by *KRT17* expression. **B**) *KRT17^+^* cells are associated with regions of subepithelial active collagen expression. **C-D**) co-expression of *COL1A1* and *KRT17* in advanced disease and **E**) presymptomatic patients with a family history of PF. Trajectory analysis of PF cells performed using RNA-velocity **F**) and Slingshot **G**) suggesting differentiation of *KRT5*^−^/*KRT17*^+^ cells from the Transitional AT2 state. **H**) RNA-ISH demonstrating *KRT17^+^* cells adjacent to *SFTPC*^+^ and *SCGB3A2*^+^ cells with low-level coexpression of multiple lineage markers in fibrotic regions of PF lung. Heatmaps depicting modules of variable genes with significant variation across pseudotime trajectories from **I**) AT2 to *KRT5*^−^/*KRT17*^+^ cells and **J**) *SCGB3A2*^+^ to *KRT5*^−^/*KRT17*^+^ cells. **K**) Smoothed expression of transcription factors with binding sites enriched for pseudotime associated genes and representative target genes across pseudotime trajectories for *KRT5*^−^/*KRT17*^+^ cells in PF and AT1 cells in controls.

Given these cells shared characteristics with airway basal cells and the distal/alveolar epithelium, the origin of these cells was unclear. To this end, we performed two trajectory analyses, based on either mRNA processing or pseudotime (see Methods) using only cells from PF lungs. Both trajectory inference methods suggest that *KRT5*^−^/*KRT17*^+^ cells are derived Transitional AT2 cells, which can arise from either AT2 or *SCGB3A2*^+^ cells (Fig. 3F-G). Consistent with this hypothesis, RNA-ISH demonstrated *KRT17^+^* cells localizing near *SCGB3A2*^+^ and *SFTPC*^+^ cells in PF lungs (Fig. 3H). To better understand the transcriptional program driving *KRT5*^−^/*KRT17*^+^ cells, we identified genes significantly associated with our pseudotime based trajectory analysis starting from both AT2 and *SCGB3A2*^+^ cells (see Methods; Fig. 3I-J, Table S6). To further characterize the regulatory program along this *KRT5*^−^/*KRT17*^+^ trajectory, we analyzed the promoters of genes associated with the trajectory for the enrichment of transcription factor binding sites (TFBS). We found that promoters of genes associated with the shift from Transitional AT2 to *KRT5*^−^/*KRT17*^+^ and stable *KRT5*^−^/*KRT17*^+^ were enriched for TFBS for the SOX family as well as binding sites for a transcriptional repressor NR1D1 (Table S7). Both *SOX4* and *SOX9* are expressed in epithelial cells, and show increased expression in PF samples (Fig. 3K). Furthermore, *NR1D1* is upregulated in AT2 and Transitional AT2s cells from control lungs, and remains near undetectable from PF samples. We then examined these key transcription factors and candidate target genes along the *KRT5*^−^/*KRT17*^+^ and AT1 trajectories and observed transcriptional changes which suggest upregulation of *SOX4* and *SOX9* combined with downregulation of *NR1D1* in AT2, *SCGB3A2*^+^, and transitional AT2 cells drives aberrant transcriptional programming leading to adoption of an ECM-producing *KRT5*^−^/*KRT17*^+^ cells (Fig. 3K, Fig. S12). Together these data provide direct evidence of an epithelial role in collagen/ECM production in PF lungs. This stands in contrast to genetic lineage tracing studies performed in mouse models of pulmonary fibrosis (*16*, *17*), likely reflecting both biological differences in the distal lung epithelium between humans and mice, mechanistic differences between human pulmonary fibrosis and the bleomycin mouse model, and the inherent limitations of lineage tracing studies using genetic Cre-lox systems and single-marker based approaches to mesenchymal cell annotation. The cytokeratin profile of this *KRT5*^−^/*KRT17*^+^ niche appears distinct from that characterizing “lineage negative epithelial progenitor” cells previously described in mouse models (*18*, *19*).

Focusing on mesenchymal cells (see Methods), we identified four discrete populations of fibroblasts in addition to smooth muscle cells and mesothelial cells (Fig. 4A). While the overall proportion of fibroblasts was elevated in PF lungs compared to controls as anticipated (Table S2), there was also a specific enrichment in fibrotic lungs of *ACTA2^+^* myofibroblasts, a *PLIN2*^+^ lipofibroblast-like group, and a previously undescribed *HAS1*^hi^ fibroblast population (Fig. S13, Table S2). Each fibroblast subtype had a unique gene expression signature (Fig. 4B,C) but there was also a collection of dysregulated genes shared across the fibroblast subpopulations (Figure 1F). Further investigation revealed the *HAS1*^hi^ population was comprised entirely of cells from IPF lungs. Analysis of genes that were upregulated in *HAS1*^hi^ cells indicated enrichment for pathways associated with cellular stress, IL4/IL13 signaling (Fig. S14) and programs previously implicated in epithelial-mesenchymal transition in other systems (*TWIST, SNAI1*) (Figure 4C, Table S4). Using RNA in-situ hybridization in matched adjacent tissue sections, *HAS1*^hi^ fibroblasts were found predominantly in subpleural regions where they colocalized with *COL1A1* expression (Figure 4D); *HAS1*^+^ cells that did not coexpress collagen were observed in subpleural regions in control lung (Fig. 4E). In contrast, we found that *PLIN2*^+^ fibroblasts were found in interstitial regions near intact alveoli in control lungs, and in PF lungs expressed lower levels of *COL1A1* than adjacent fibroblasts (Fig. 4F,G). Myofibroblasts identified by coexpression of *MYLK*^+^/*COL1A1*^+^ localized primarily to subepithelial regions near airways (Figure 4H,I). Together, these data suggest a global dysregulation of fibroblasts occurs in PF along with expansion of specific pathogenic subtypes in spatially distinct niches driving pathologic ECM deposition and expansion.

**Figure 4.**
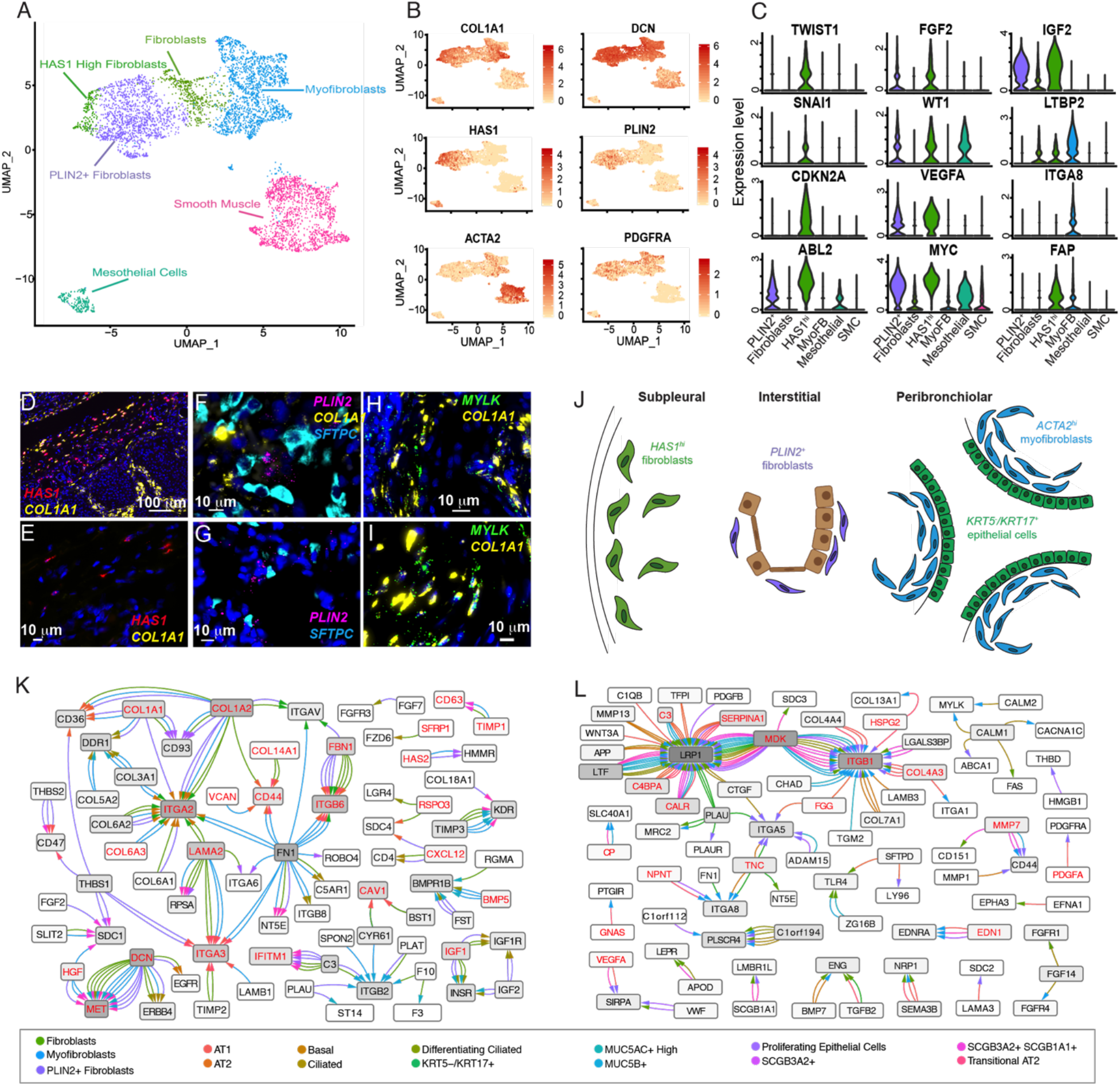
Characterization of mesenchymal stromal cell types in PF lung. **A**) Identification of 4 distinct fibroblast populations **B**) characterized by expression of classical and novel markers including *HAS1*. **C**) Expression of discrete activation programs among fibroblast subtypes. **D**) *HAS1^hi^* fibroblast coexpressed *COL1A1* in subpleural regions of PF lungs, while **E**) *HAS1^hi^* cells lacking collagen expression were founding in analogous regions of control lungs. **F**) *PLIN2^+^* fibroblasts were diffusely located in interstitial regions and expressed low-levels of *COL1A1* in PF lungs. **G**) In control lungs, *PLIN2^+^* fibroblasts were located near *SFTPC*^+^ AT2 cells. **H-I**) *MYLK*^+^ myofibroblasts were locating in regions morphologically resembling fibroblastic foci in subepithelial regions. **J**) Schematic summarizing discrete fibroblast populations identified in PF lungs. **K**) Interactome based on the coexpression of fibroblast ligands and epithelial receptors and **L**) epithelial ligands and fibroblast receptors; edges are colored by the ligand-expressing cell type; arrowheads are colored by the receptor-expressing cell type; red lettering indicates differential expression between PF and control lungs.

In order to determine potential mediators of pathologic mesenchymal-epithelial communication, we performed an interactome-based analysis identifying putative ligand:receptor binding pairs in epithelial and mesenchymal cells (see Methods) and constructed interaction networks for mesenchymal-driven (Figure 4K) and epithelial-driven (Figure 4L) signaling (Fig. S15, Table S8). Both networks were significantly enriched for genes differentially expressed between PF and control lungs (see Methods). This analysis implicated matrix-driven signaling through integrin receptors as the central mechanisms through which fibroblast lineages interact with epithelial cells in PF lungs. In contrast, a more complex network involving multiple growth factors, cytokines and chemokines signaling through integrins, Wnt-coreceptors, and other pathways was observed for epithelial-driven signaling.

Together, these data provide substantial insight into the complexity, heterogeneity, and plasticity of the peripheral lung in human disease, building upon molecular atlasing efforts in the diseased (*20*, *21*) and healthy (*22*) lung. This high-resolution overview identifies the genes, pathways and programs that characterize pathologic lung remodeling in pulmonary fibrosis. Future studies investigating the origin, behavior and function of these novel cell types, subtypes, states and pathologic gene expression programs should provide additional insights into the foundational mechanisms regulating homeostasis and disease in the human lung.

## Data and code accessibility

Raw and processed 10X genomics data can be found on GEO using accession number: GSE135893. The code used to analyse the data can be found at https://github.com/tgen/banovichlab/.

## Acknowledgements

This work was supported by NIH/NHLBI R01HL145372(JAK/NEB), K08HL130595 (JAK), P01HL92870(TSB), the Doris Duke Charitable Foundation (JAK), R01HL126176 (LBW), K08HL136888(CMS), K08HL143051 (JMMS), T32HL094296 (NIW), Department of Veterans Affairs (BWR), Boehringer Ingelheim Pharmaceuticals, Inc (BIPI). BIPI had no role in the design, analysis or interpretation of the results in this study; BIPI was given the opportunity to review the manuscript for medical and scientific accuracy as it relates to BIPI substances, as well as intellectual property considerations. Vanderbilt Cell Imaging Shared Resource is supported by NIH grants CA68485, DK20593, DK58404, DK59637 and EY08126. We would like to thank 10X genomics, particularly Jill Herschleb, for their help and support of early protocol optimizations. We also want to thank the Chan Zuckerberg Initiative, who supported early efforts in the lab to optimize scRNA-seq on the lung. Finally, we thank Tennessee Donor Services and the Donor Network of Arizona, and the patients and families who generously donated tissue samples to make these studies possible.

## Author Contributions

NEB and JAK conceived and designed the study; ACH, NIW, CLC, LP, MIC processed biospecimens and prepared single-cell suspensions; LP, NEB, LR, JR prepared sequencing libraries and performed NGS; CJT, CJ, JMSS, JAK performed in-situ hybridization and microscopy; JEL, MB, LBW, CMS, RB, RW procured tissue specimens; LP, GD, LW collected and analyzed clinical data; ACH, AJG, LTB, SLY, NEB, JAK performed quality checks, data integration and computational analyses; ACH, AJG, LTB, NIW, BWR, APS, WJM, SBM, TSB, NEB, JAK analyzed and interpreted scRNA-seq data; ACH, AJG, LTB, NEB, JAK wrote and revised manuscript

## Supplementary Materials

Materials and Methods

Figures S1-S15

Tables S1-S8

## Notes

https://github.com/tgen/banovichlab/

