## Supplementary material for "Single-cell RNA-sequencing reveals profibrotic roles of distinct epithelial and mesenchymal lineages in pulmonary fibrosis": Supplementary information.pdf

**This PDF file contains**

Materials and Methods
Figures S1-S15
Tables S1-S8

**Materials and Methods**

The code for genomic analyses in this paper is available at https://github.com/tgen/banovichlab/

*Subjects and samples*: PF tissue samples were obtained from lungs removed at the time of lung transplantation at two lung transplant centers (VUMC and NTI). Nonfibrotic control tissue samples were obtained from lungs declined for organ donation. For PF lungs, diagnoses were determined according to ATS/ERS consensus criteria (*1*). All studies were approved by the local Institutional Review Boards (Vanderbilt IRB #'s 060165, 171657, Western IRB # 20181836).

*Tissue Processing*: Biopsies of multiple regions from each lung sample were digested in an enzymatic cocktail (collagenase I/dispase II 1 μg/ml tissue or Miltenyi Multi-Tissue Dissociation Kit) using a gentleMACS Octo Dissociator (Miltenyi, Inc). Adjacent tissue was fixed in 10% formalin for 24-72 hours and used for tissue localization studies. Tissue lysates were serially filtered through sterile gauze, 100 um then 40 um sterile filters (Fischer). Single cell suspensions then underwent cell sorting using serial columns (Miltenyi Microbeads, CD235a, CD45) at VUMC or FACS at TGen. CD45- and C45+ populations were mixed 2:1 in samples processed at VUMC were used as input for generation of scRNA-seq libraries. At TGen, Calcien-AM was used to stain live cells and 10,000-15,000 total live cells were sorted directly into the 10X reaction buffer and transferred immediately to the 10X 5' chip A (10X Genomics).

*scRNA-seq library preparation and next-generation sequencing:* ScRNA-seq libraries were generated using the 10X Chromium platform 3' v2 or 5' library preparation kits (10X Genomics) following the manufacturer's recommendations and targeting 5,000-10,000 cells per sample. Next generation sequencing was performed on an Illumina Novaseq 6000 or HiSeq 4000. Reads with read quality less than 30 were filtered out and CellRanger Count v3.0.2 (10X Genomics) was used to align reads onto GRCh38 reference genome.

*In-situ hybridization and microscopy*: RNAscope performed according to manufacturer's instructions using the following probes and reagents: SFTPC-C1 (Cat# 452561), COL1A1-C2 (Cat# 401891-C2), K17-C3 (Cat# 463661-C3), K5-C2 (Cat# 310241-C2), HAS1-C1 (Cat# 483251), MYLK-C3 (Cat# 533471-C3), PDGFRA-C4 (Cat# 604481-C4), Hs-SCGB3A2-C1 (Cat# 549951), Multiplex v2 kit (Cat# 323100). Briefly, tissue was fixed in 10% neutral buffered formalin for at 4C for 72 hours then embedded and sectioned. Slides were deparaffinized and allowed to completely dry. Endogenous peroxidase activity was quenched with hydrogen peroxide for 10 min at room temperature. Target retrieval was performed in ACDbio RNAscope 1x Target Retrieval Reagent at 99-102 °C for 15 min. A hydrophobic barrier was drawn around the tissue with Immedge Pap Pen (Vector Labs, Cat# H-4000) and allowed to dry overnight. We applied RNAscope Protease Plus for 15 minutes at RT then proceeded to run the RNAscope assay. We hybridized the probes, applied RNAscope signal amplifiers, and labeled probes accordingly. Tissue was exposed to DAPI for 30 seconds at room temperature and then mounted in Prolong Gold and allowed to dry overnight at room temperature. Immunofluoresence images were acquired using a Keyence BZ-X710 with BZ-X Viewer software and/or an automated TiE inverted fluorescence microscope platform equipped with an encoded motorized stage and Plan Apo 60x 1.40 NA objective (Nikon Instruments, Inc.) and additionally outfitted with a Yokogawa X1 spinning disk head and Andor DU-897 EM-CCD. Lasers utilized for excitation included 405, 488, 561, and 647 nm lines. Image stitching was performed using the BZ-X Analyzer package. Emission filters were 455/50, 525/36, 641/75, and 700/74 (peak/bandwidth), respectively. NIS-Elements software (Nikon Instruments, Inc.) was utilized for acquisition.

*Dimensionality reduction, clustering and visualization*: Seurat v3 was used to perform dimensionality reduction, clustering and visualization for the scRNAseq data (*2*, *3*). Individual sample output files from CellRanger Count were read into Seurat v3 to generate a unique molecular identified (UMI) count matrix that was used to create a Seurat object containing a count matrix and analysis. All Seurat objects were combined into a merged dataset and percentage of mitochondrial genes were calculated for each sample in the merged object. Cells containing less than 1,000 identified genes and more than 25% percentage of reads arising from mitochondrial genes removed. (Fig S1). SCTransform with default parameters was used to normalize and scale the data, and dimensionality reduction was performed using PCA on the top 3,000 most variable genes. To determine the optimal number of Principal Components (PCs) for UMAP visualizations and to avoid overfitting, we attempted to identify an optimal number of PCs that kept the relative distance between points on UMAP-1 and UMAP-2 stable - i.e. the UMAP plot remained stable between across a range of PCs used to generate the UMAP plot. To this end, we used Mantel randtest to calculate correlation between relative location of points on the UMAP plot between two adjacent number of included PCs for both UMAP-1 or UMAP-2. Correlation coefficient values were plotted and we manually selected PC ranges where both UMAP-1 and UMAP-2 plateaued. The smallest PC number within the plateau was chosen for input into the final UMAP.

*Cell type annotation and doublet removal*: Markers specific for major cell types: *PTPRC*$^{+}$ (Immune cells), *EPCAM*$^{+}$ (Epithelial cells), *PECAM1*$^{+}$*/PTPRC*$^{-}$ (Endothelial cells) and *PTPRC*$^{-}$*/EPCAM*$^{-}$*/PECAM1*$^{-}$ (Mesenchymal cells) were used to split Seurat clusters into four sub groups (Fig S2). Each sub group object underwent the same dimensionality reduction, clustering, and visualization approach as described above. Each sub group object was then further split into clusters and manually annotated with known cell type markers (Table S3). Doublet cells were identified manually as expressing markers for different cell types and the final object was created by merging all four annotated, doublet-removed sub groups.

*Differential expression analysis:* To identify differentially expression genes (DEGs) between cell types, we used a negative binomial model as implemented in the Seurat FindMarkers function comparing each individual cell type to all other cells within the major cell type cluster (Immune, Epithelial, Endothelial, and Mesenchymal). Genes were considered differentially expressed if the adjusted P value was lower than 0.01 (Table S4). To identify genes that were differentially expressed between PF and control lungs, we took each cell type independently - for all cell types with a minimum of 50 cells in both PF and control lungs - and used the negative binomial model to test for differences in expression. Genes were considered differentially expressed if the adjusted P value was less than 0.01 (Fig. 1D, Table S4).

*Cell trajectory analysis*: Two methods were used to perform trajectory analysis: **A**) We used RNA velocity (*23*), a method based on mRNA processing to calculate cell trajectories The run10x function was used to annotate spliced, unspliced, and spanning reads from our scRNA-seq data (*23*, *24*). Genes with an average expression count of less than 0.1 for spliced reads, 0.05 for unspliced and 0.001 for spanning reads were removed. After applying QC filtering, gene relative velocity estimates were calculated using a fit quantile of 0.01 and kCells of 50. **B**) The R package Slingshot (*11*) was used to perform a pseudotime based cell trajectory analysis. The start and end cluster assignment was based on the results from RNA velocity analysis. slingshot wrapper function was performed with the UMAP dimensionality reduction and cluster labels as in Seurat objects to identify the trajectory.

*Identifying genes associated with trajectory analysis:* To identify genes along the course of the trajectory, a general additive model (GAM) was used to regress each gene on the pseudotime variable. The top 400 significant genes were chosen for heatmap based on loess GAM p-values and heatmap was plotted using the function plotHeatmap in the R package clusterExperiment (*25*). Loess plot for individual gene of interest was generated using ggplot2 geom_smooth function (smoothing method: “loess”) on the trajectory variable timeline resulted from the slingshot wrapper function above.

*Transcription Factor Binding Site Analysis:* for motif analysis, findMotifs.pl (package HOMER v4.10) (*26*) was used on promoter sequences (700bp upstream and 100bp downstream of TSS) of genes in the trajectory heatmap (Figure 3.I, Table S.6) using HOMER pre_built human promoter database and default parameters. To identify SOX and NR1D1 motif location, HOMER annotatePeaks.pl was performed on promoter regions of the targeted genes using the motif files generated from findMotifs.pl function.

*Pathway Enrichment Analysis*: Pathway enrichment was performed using PANTHER Pathways through the WebGestalt 2019 (*27*). Differentially expressed genes (FDR <0.05) with an absolute increase in proportion of cells expressing a given gene of >0.1 and log fold-change >0.4 (*HAS1$^{hi}$* vs. other mesenchymal cells) with were selected as input for enrichment analysis.

*Interactome Analysis:* The scaled gene expression matrix for diseased cells from a Seurat sub-group object containing epithelial cells and mesenchymal cells (excluding HAS1$^{hi}$ cells, which localized to subpleural regions and were not in spatial proximity with non-mesothelial epithelial populations) was filtered to contain the top 20% highly expressed genes for each cell type using the iTALK function rawParse (*28*). These top genes were used as input for the iTALK FindLR function to find ligand-receptor (LR) pairs between all cell types. Hundreds of LR pairs were found between epithelial cells and mesenchymal cells (Table S8). Next, we tested if highly co-expressed LR pairs (top 20%) were enriched for differentially expressed genes between ILD and control samples. First, we generated an empirical null distribution for each cell type interaction (excluding PLIN2+ fibroblasts and KRT5-/KRT17+ epithelial cells, which did not contain sufficient numbers of control cells for differential expression testing) by randomly sampling a matched number (Table S8) of LR pairs from all LR pairs identified for that interaction from the iTALK database 1,000 times and identifying the number of significantly differentially expressed genes within each iteration. P-values were calculated as the proportion of permutations that exceeded the number of differentially expressed genes within the top 20% most highly co-expressed pairs. For highly co-expressed ligands from mesenchymal cells and corresponding receptors in epithelial cells, significant enrichment (p-value < 0.05) was found for LR pairs between fibroblasts and each epithelial cell type, as well as between myofibroblasts and each epithelial cell type, excluding ciliated cells (p = 0.083). For highly co-expressed ligands from epithelial cells and corresponding receptors in mesenchymal cells, significant enrichment was found for LR pairs between fibroblasts and each epithelial cell type excluding ciliated (p = 0.071), differentiating ciliated (p = 0.53), and proliferating epithelial cells (p = 0.168), as well as between myofibroblasts and each epithelial cell type, excluding ciliated (p = 0.068) and differentiating ciliated cells (p = 0.179). For Figure 4K-L, the top 5 most highly co-expressed pairs (ranked by the product of mean ligand expression and mean receptor expression) for each epithelial cell - mesenchymal cell interaction were visualized using Cytoscape (*29*). Differentially expressed genes within the visualized LR pairs are denoted by red text.

### Supplementary Figures

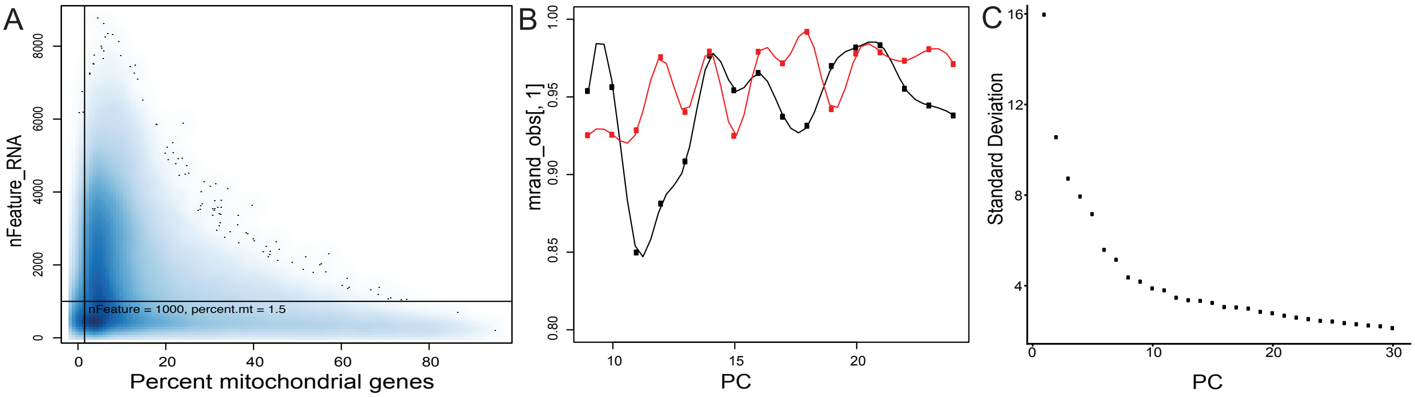

**Figure S1.** Density plot demonstrating distribution of percentage of mitochondrial genes and nFeature_RNA in all scRNA-seq samples (A). Correlation coefficient values (mrand_obs) between two adjacent UMAP-1 and UMAP-2 (B) and Elbow Plot showing standard deviation between adjacent PC for 30 PCs (C).

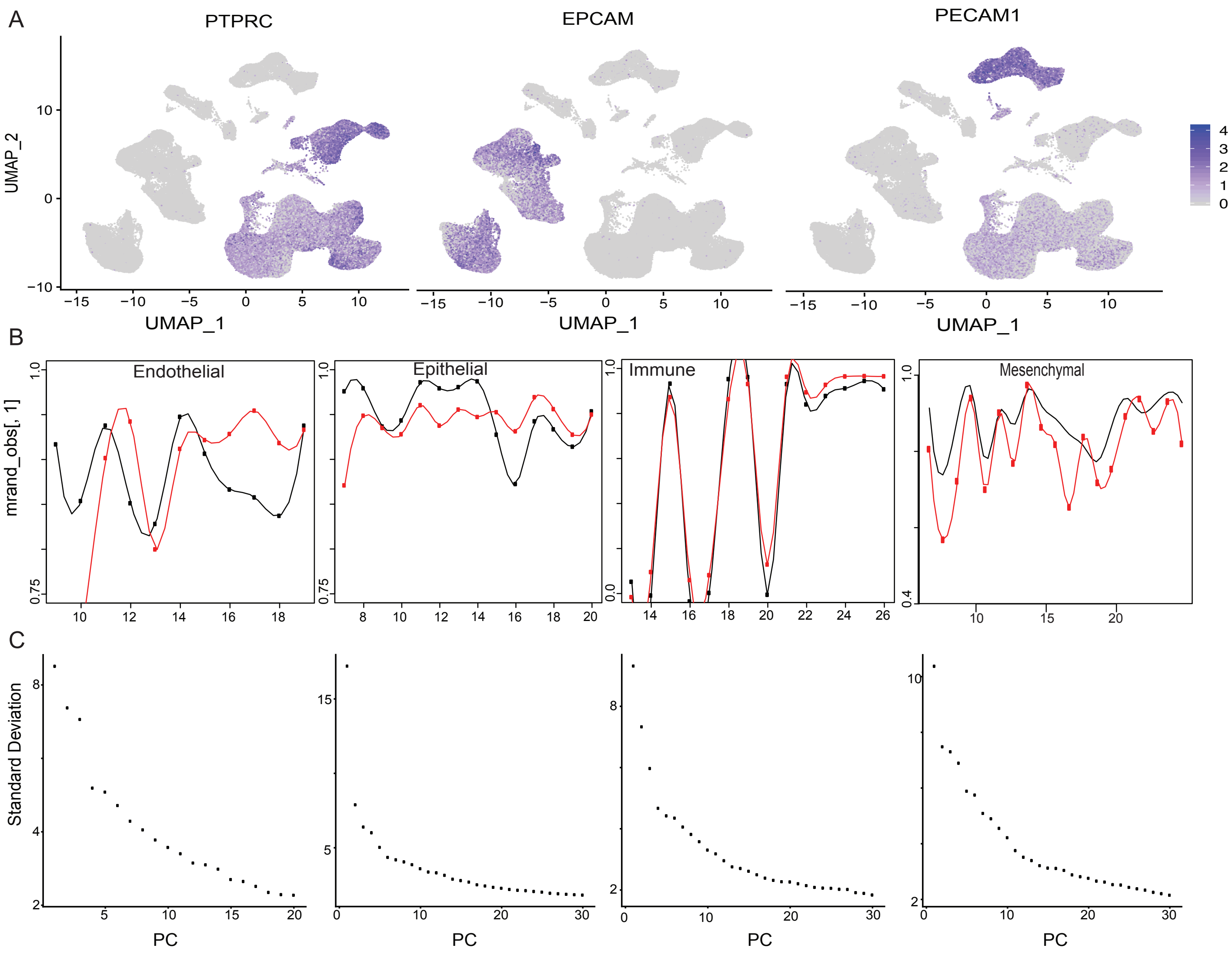

**Figure S2.** Feature Plots showed expression of cell type specific markers (*PTPRC*, *EPCAM* and *PECAM1*) on Seurat clusters (A). Correlation coefficient values (mrand_obs) between two adjacent UMAP-1 and UMAP-2 (B) and Elbow Plot showing standard deviation between adjacent PC for 30 PCs (C).

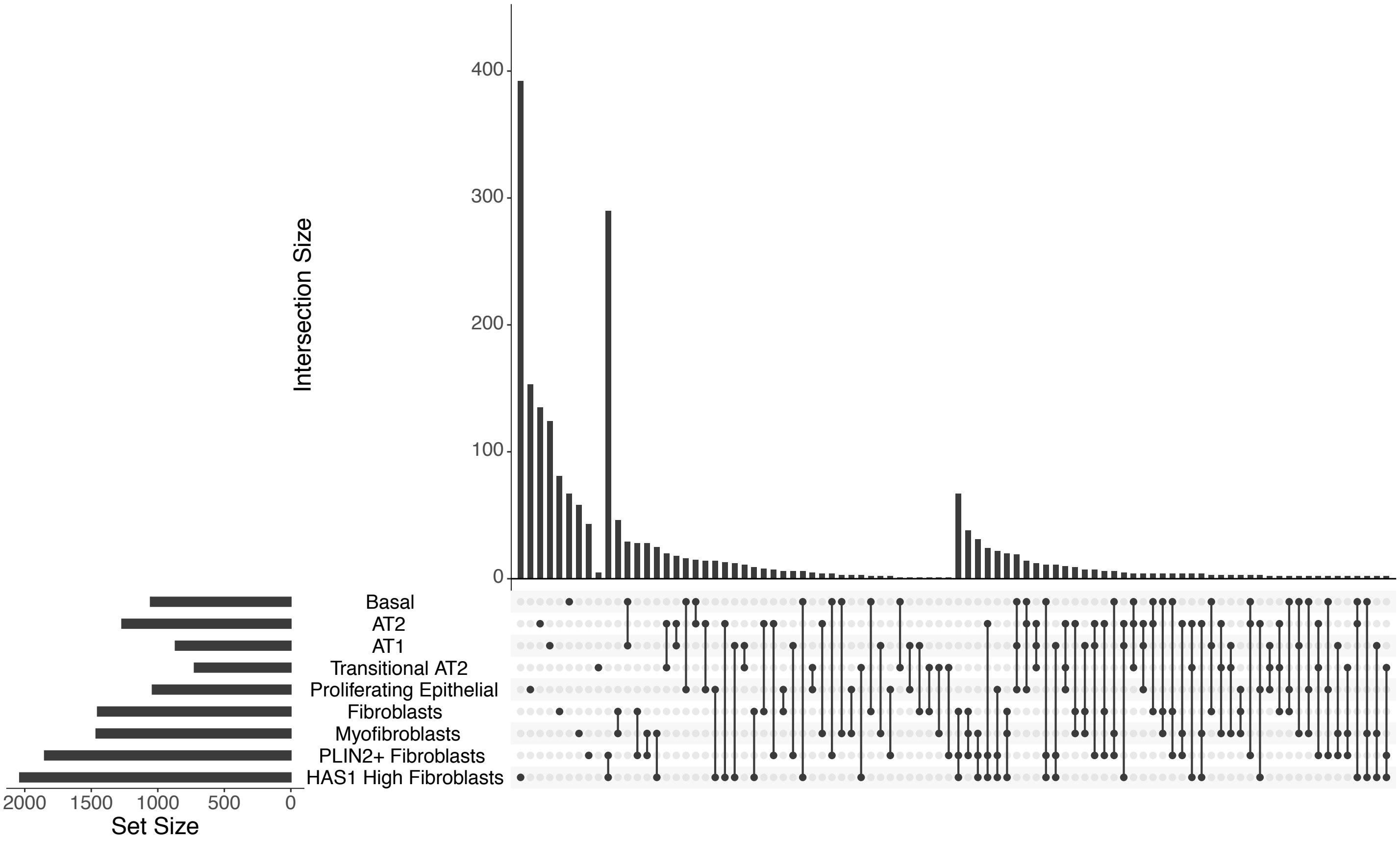

**Figure S3**. Shared DEGs between KRT5-/KRT17+ compared to epithelial and fibroblast cell types among IPF samples

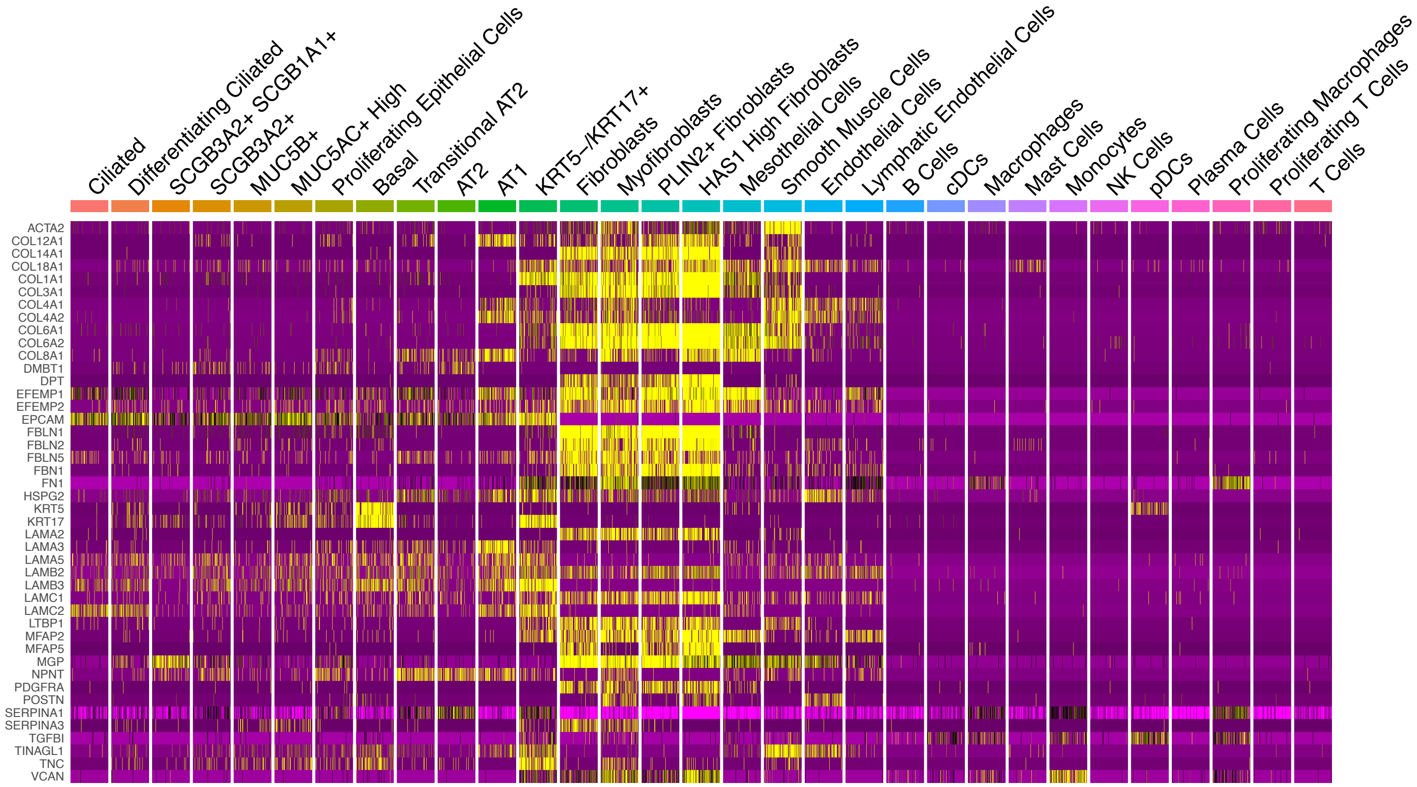

**Figure S4**. Heatmap demonstrating relative expression of IPF-enciched ECM components including all cell-types (expanded from Figure 1F).

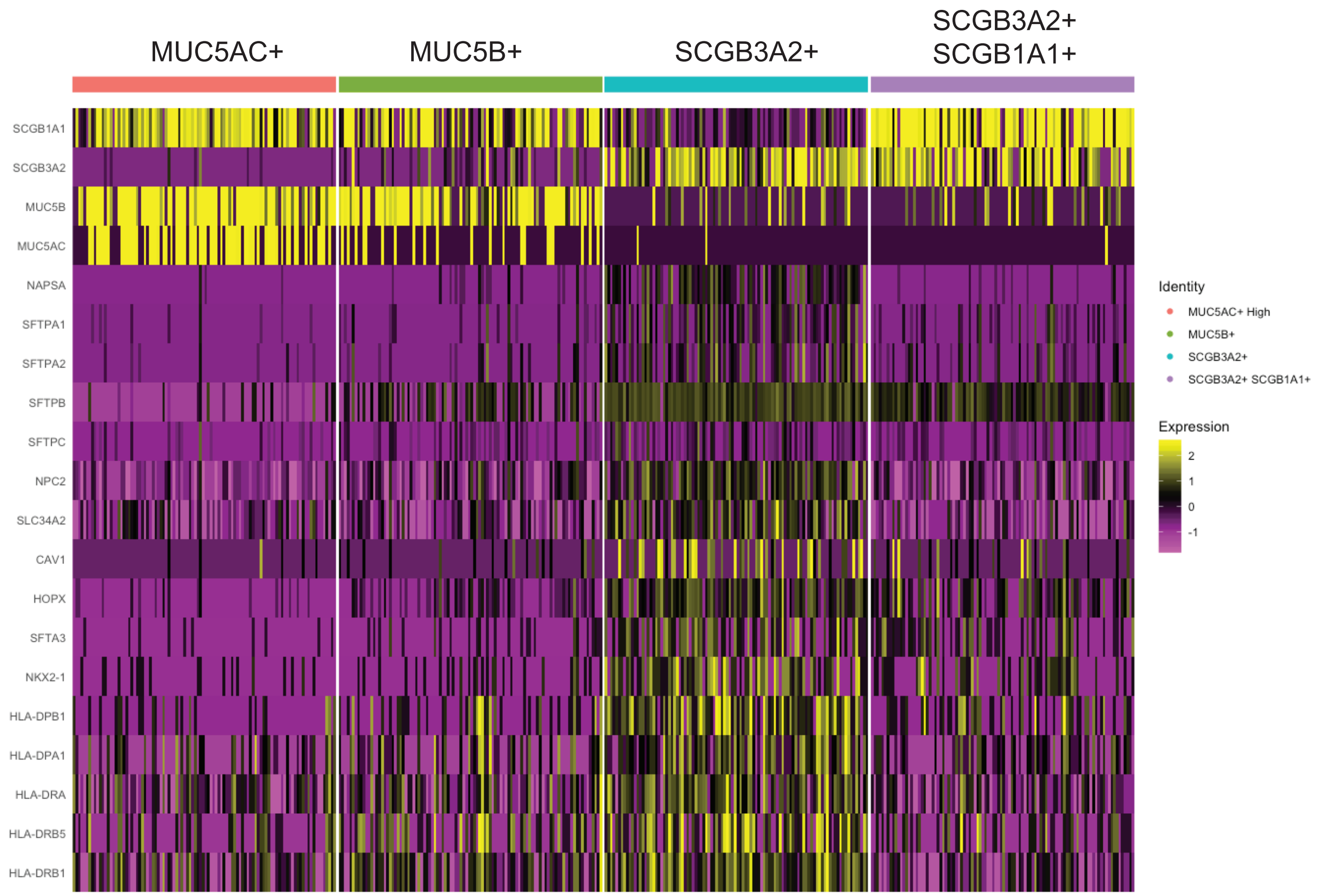

**Figure S5.** Heatmap demonstrating selected markers differing significantly (FDR<0.05, log FC > 0.4) between *SCGB3A2+* and other secretory cell subtypes.

A

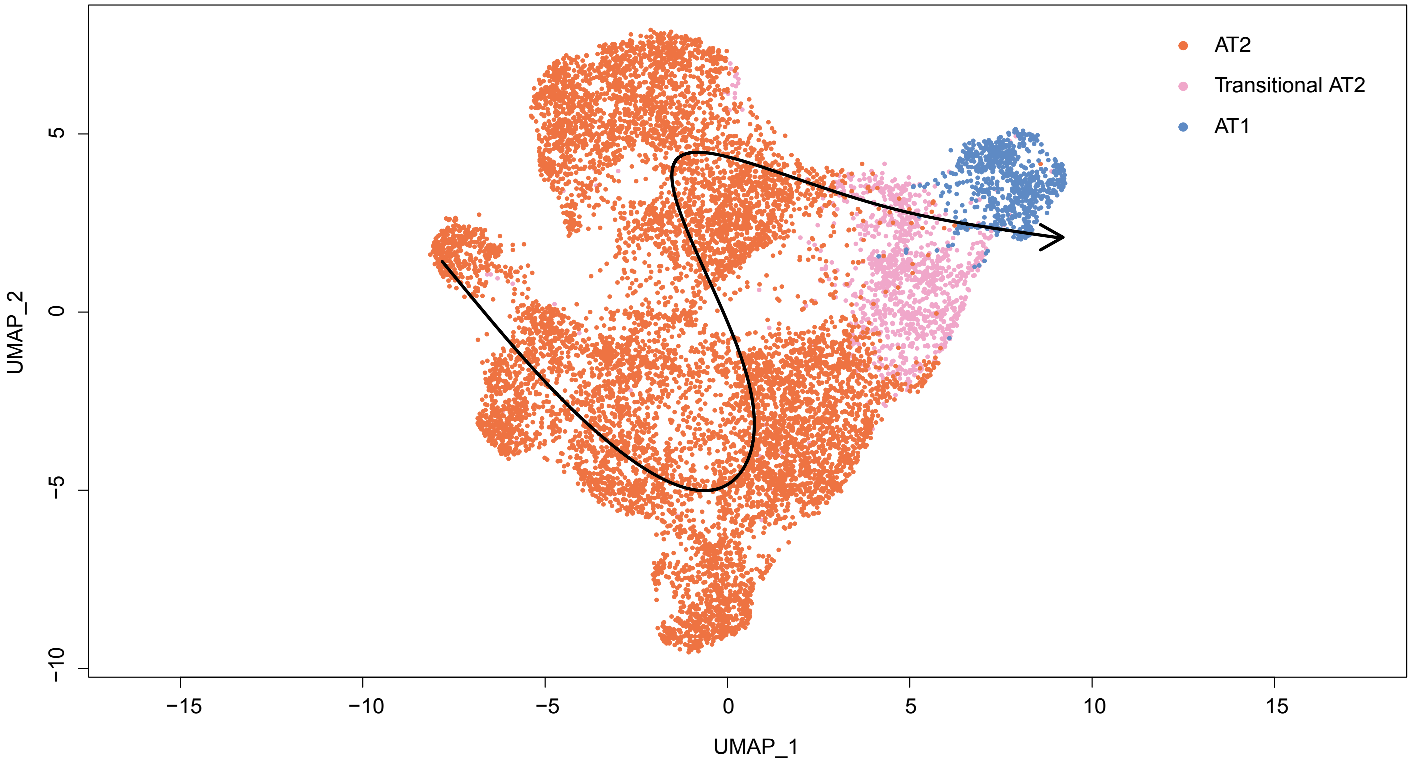

B

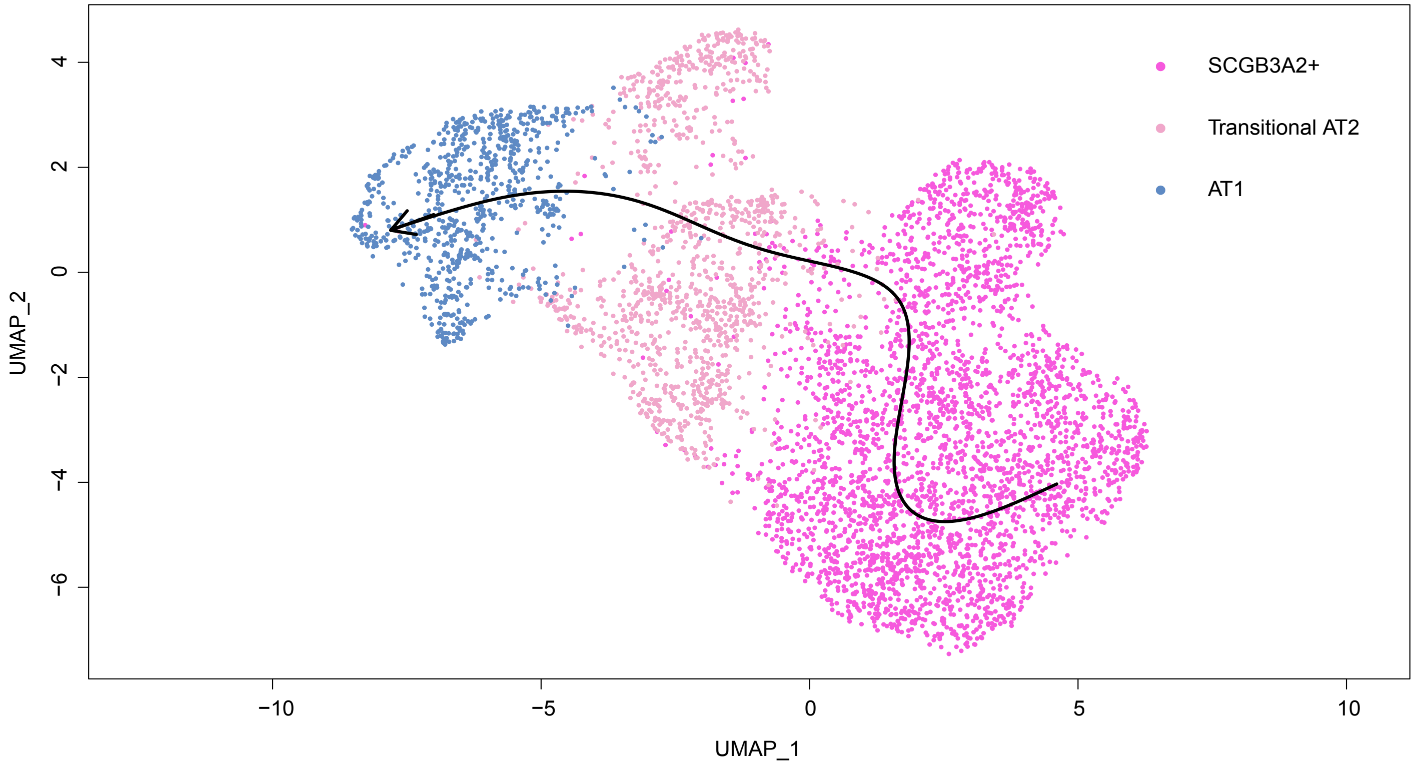

**Figure S6**. Slingshot trajectory analysis demonstrating the transition from AT2 (A) or SCGB3A2+ (B) into transitional AT2 and AT1 cell types in fibrotic and control lungs (used for the loess plot in Figure 2G)

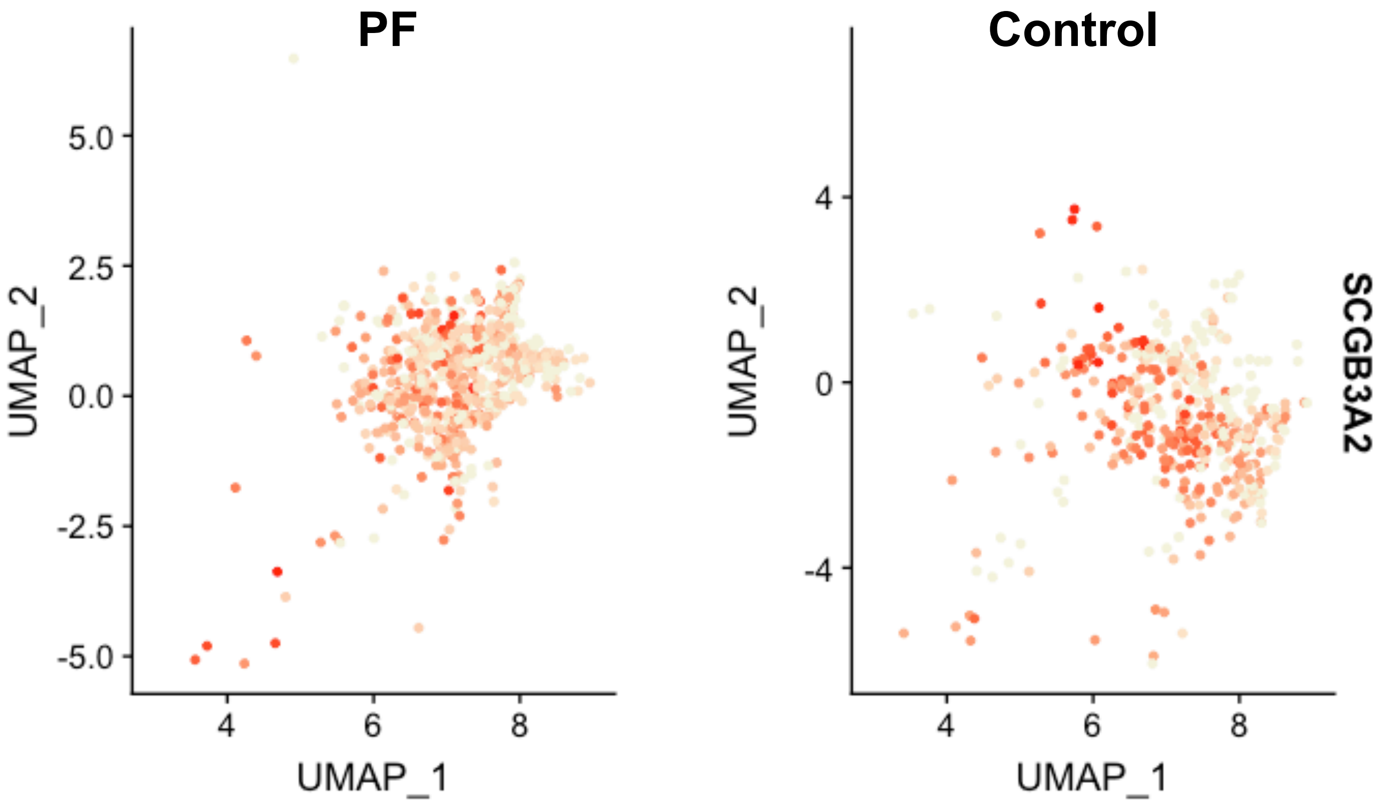

**Figure S7**. Expression of SCGB3A2 in transitional AT2 cells in PF and control Transitional AT2 cells.

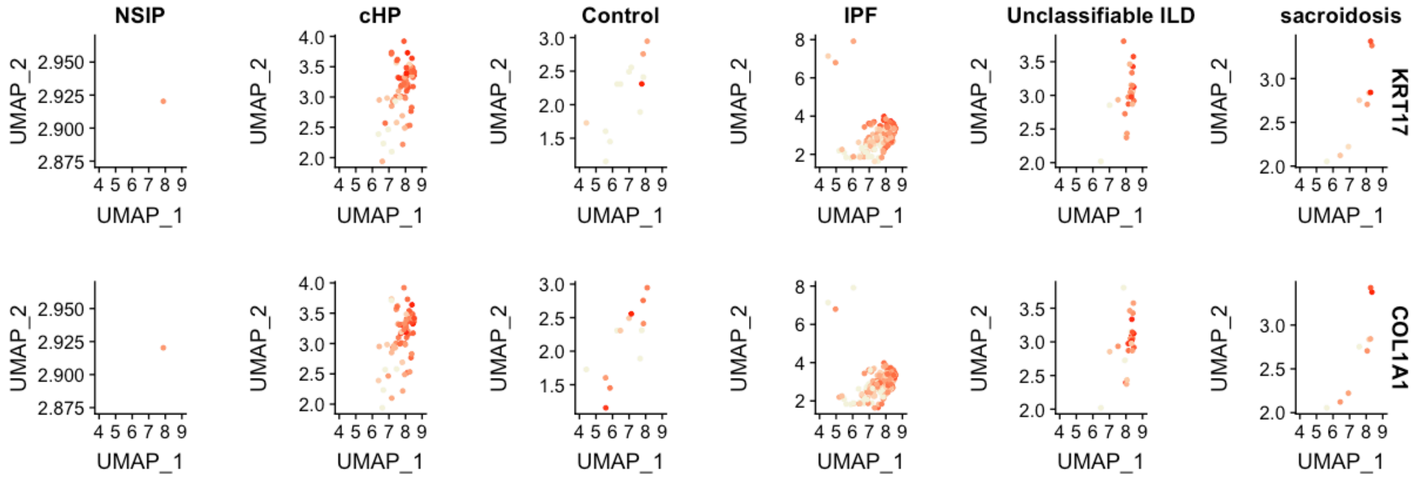

**Figure S8**. Expression of *COL1A1* and *KRT17* in *KRT5-/KRT17+* cells comparing different histopathologic patterns of PF.

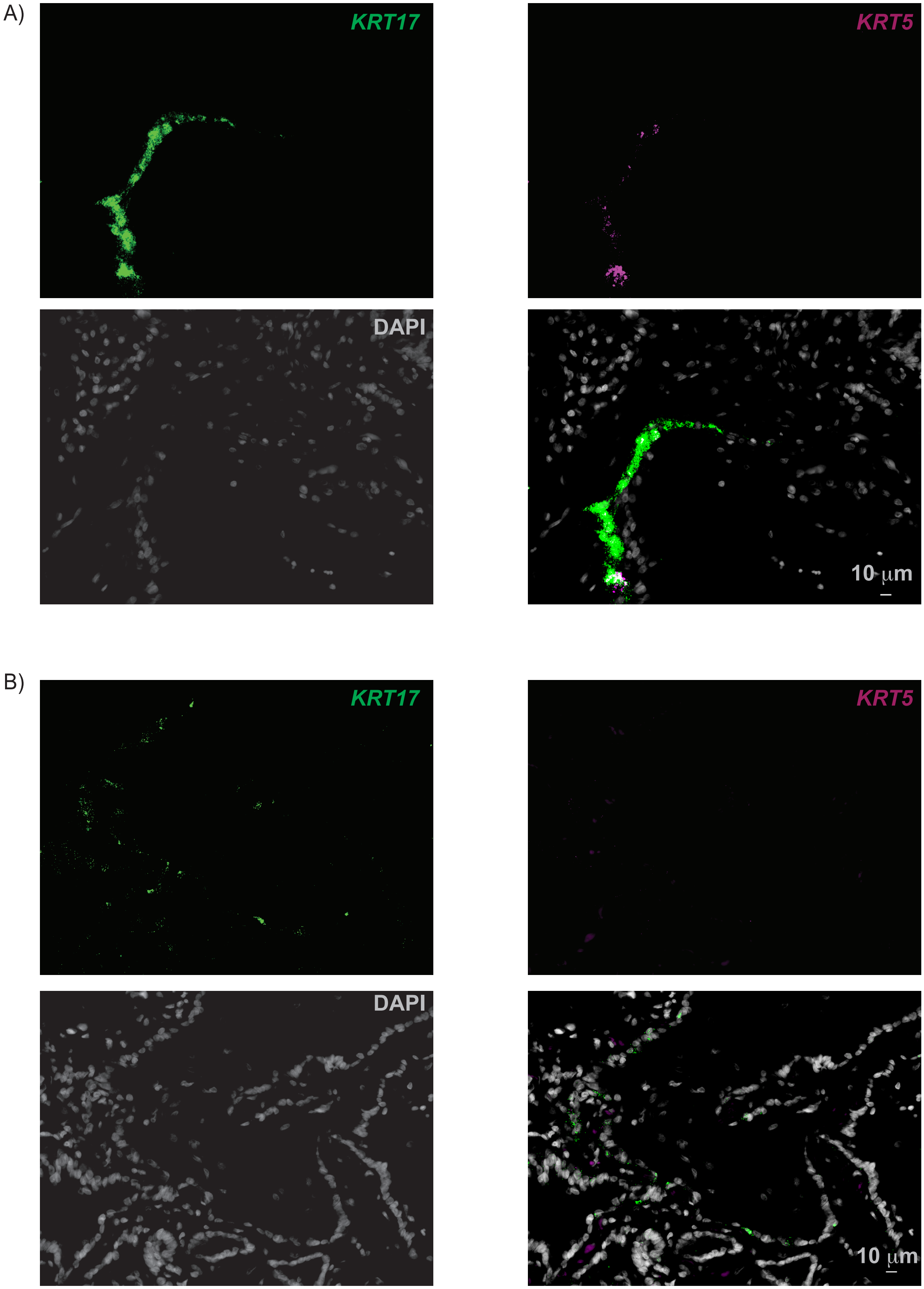

**Figure S9**. Multiplexed RNA-ISH for *KRT5* and *KRT17* in A) large airways and B) peripheral lung from PF lungs.

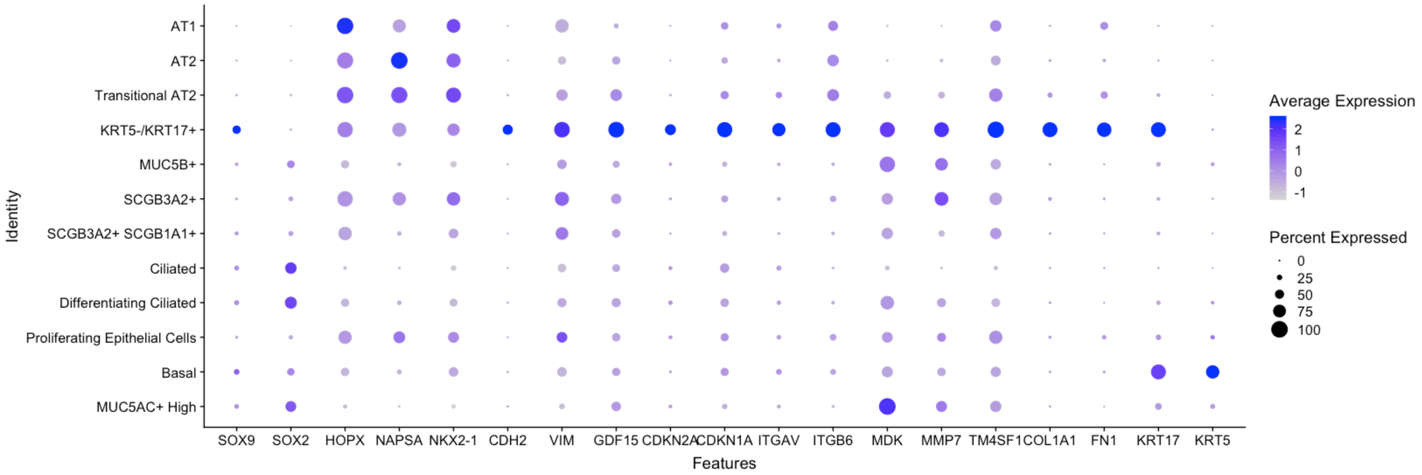

**Figure S10**. Relative expression of selected top *KRT5-/KRT17+* markers compared to other epithelial cell types.

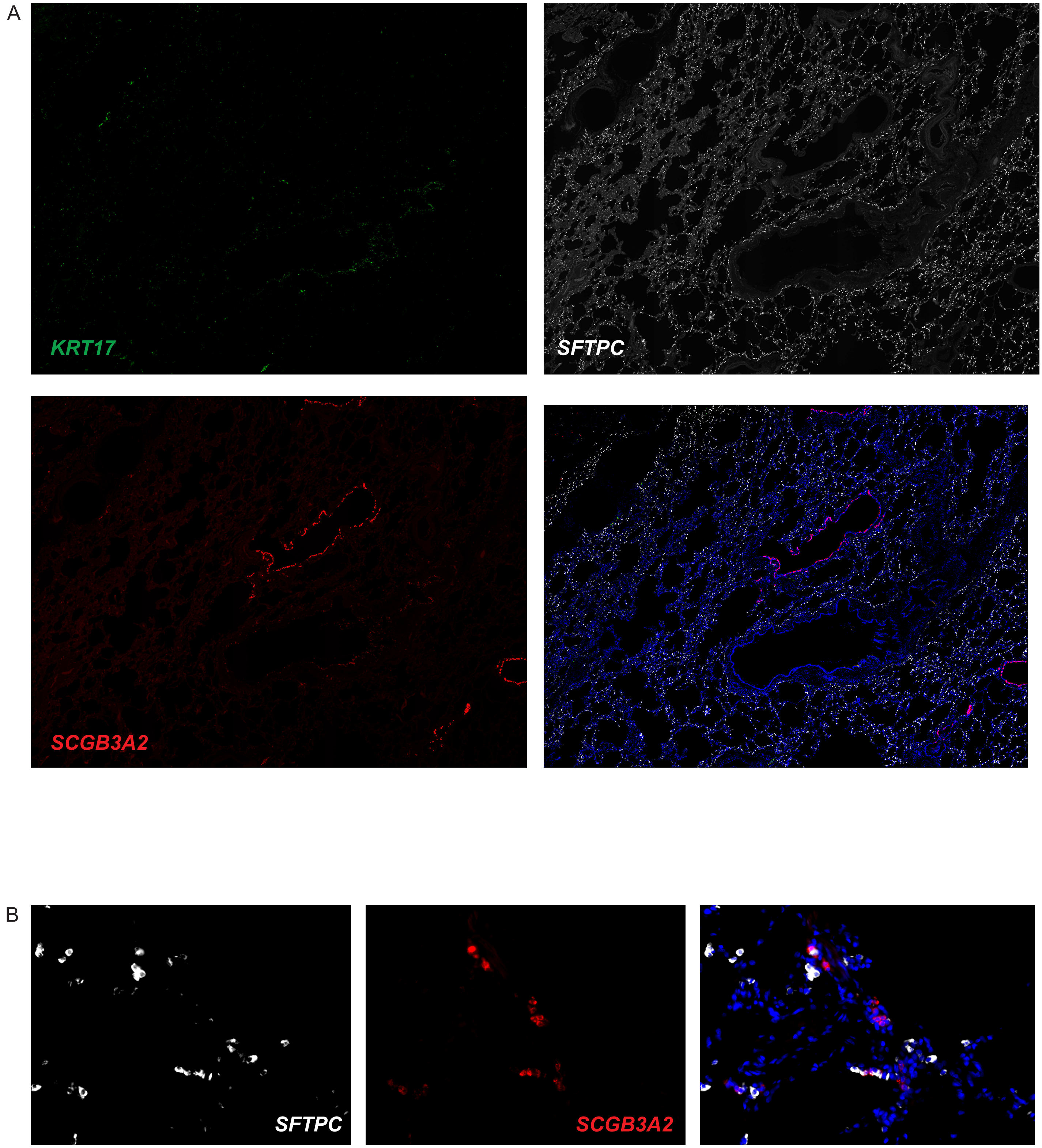

**Figure S11**. A) Multiplexed RNA-ISH for *SCGB3A2*, *SFTPC* and *KRT17* in control lung (40x original magnification). B) Identification of rare *SCGB3A2*+/*SFTPC*+ dual-positive cells in control lungs (original magnification 400x).

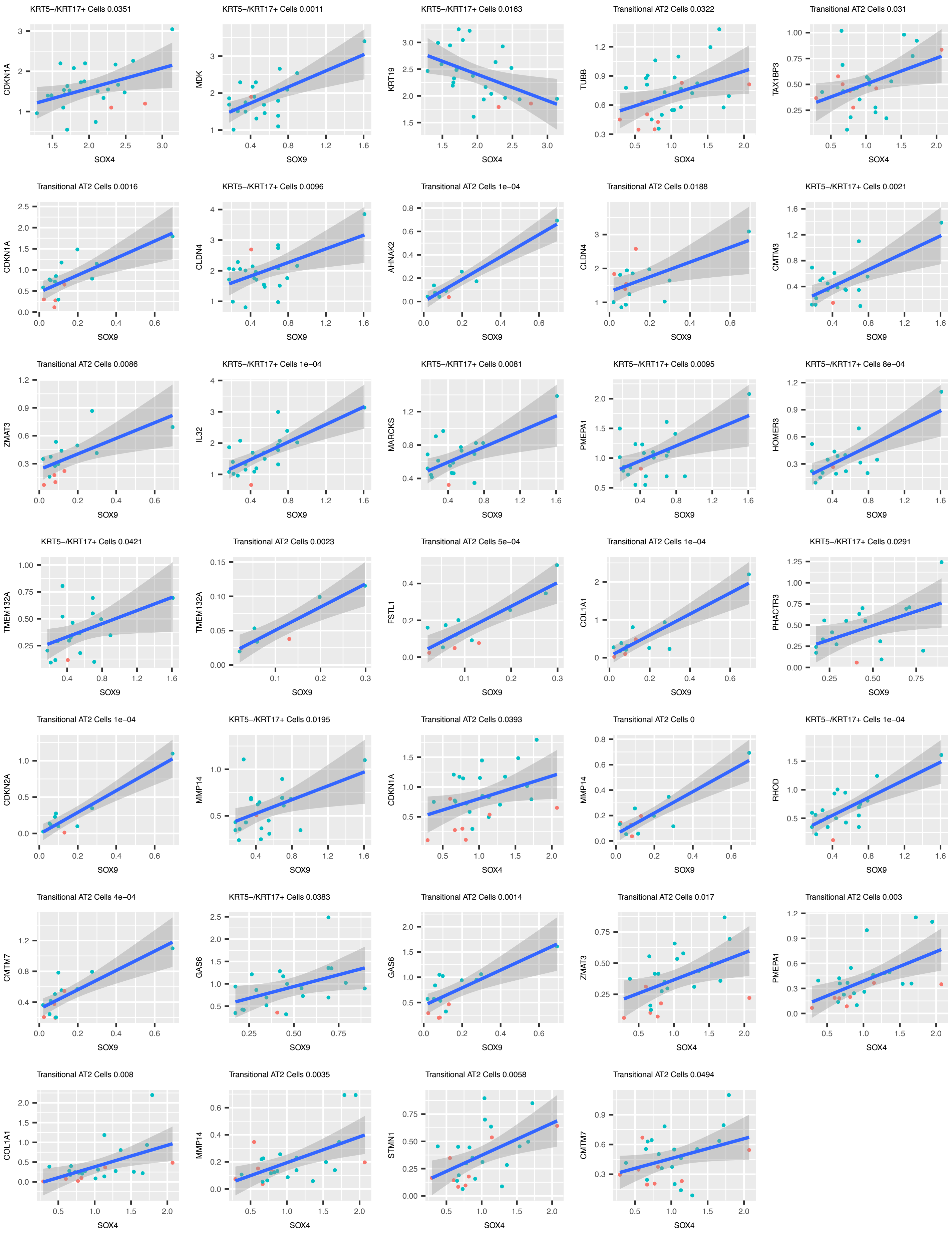
KRT5-/KRT17+ Cells 0.0351
CDKN1A
SOX4
KRT5-/KRT17+ Cells 0.0011
MDK
SOX9
KRT5-/KRT17+ Cells 0.0163
KRT19
SOX4
Transitional AT2 Cells 0.0322
TUBB
SOX4
Transitional AT2 Cells 0.031
TAX1BP3
SOX4
Transitional AT2 Cells 0.0016
CDKN1A
SOX9
KRT5-/KRT17+ Cells 0.0096
CLDN4
SOX9
Transitional AT2 Cells 1e-04
AHNAK2
SOX9
Transitional AT2 Cells 0.0188
CLDN4
SOX9
KRT5-/KRT17+ Cells 0.0021
CMTM3
SOX9
Transitional AT2 Cells 0.0086
ZMAT3
SOX9
KRT5-/KRT17+ Cells 1e-04
IL32
SOX9
KRT5-/KRT17+ Cells 0.0081
MARCKS
SOX9
KRT5-/KRT17+ Cells 0.0095
PMEPA1
SOX9
KRT5-/KRT17+ Cells 8e-04
HOMER3
SOX9
KRT5-/KRT17+ Cells 0.0421
TMEM132A
SOX9
Transitional AT2 Cells 0.0023
TMEM132A
SOX9
Transitional AT2 Cells 5e-04
FSTL1
SOX9
Transitional AT2 Cells 1e-04
COL1A1
SOX9
KRT5-/KRT17+ Cells 0.0291
PHACTR3
SOX9
Transitional AT2 Cells 1e-04
CDKN2A
SOX9
KRT5-/KRT17+ Cells 0.0195
MMP14
SOX9
Transitional AT2 Cells 0.0393
CDKN1A
SOX4
Transitional AT2 Cells 0
MMP14
SOX9
KRT5-/KRT17+ Cells 1e-04
RHOD
SOX9
Transitional AT2 Cells 4e-04
CMTM7
SOX9
KRT5-/KRT17+ Cells 0.0383
GAS6
SOX9
Transitional AT2 Cells 0.0014
GAS6
SOX9
Transitional AT2 Cells 0.017
ZMAT3
SOX4
Transitional AT2 Cells 0.003
PMEPA1
SOX4
Transitional AT2 Cells 0.008
COL1A1
SOX4
Transitional AT2 Cells 0.0035
MMP14
SOX4
Transitional AT2 Cells 0.0058
STMN1
SOX4
Transitional AT2 Cells 0.0494
CMTM7
SOX4

**Figure S12.** Significant correlation between the transcription factors SOX4 and SOX9 and genes containing motifs for those transcription factors in the Transitional AT2 and KRT5-/KRT17+ cells across individuals.

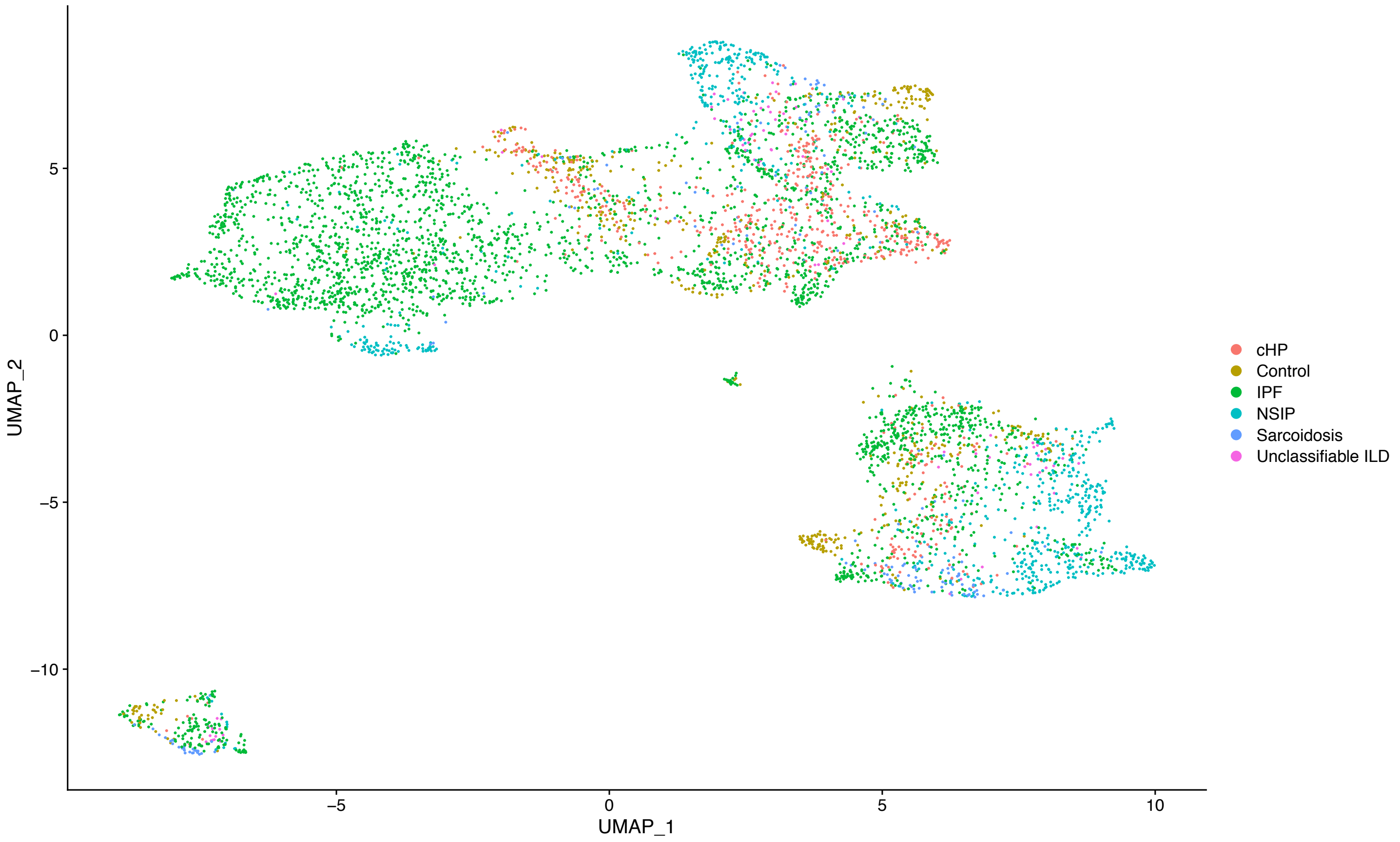

**Figure S13**. Mesenchymal cell types annotated by histopathologic pattern of PF.

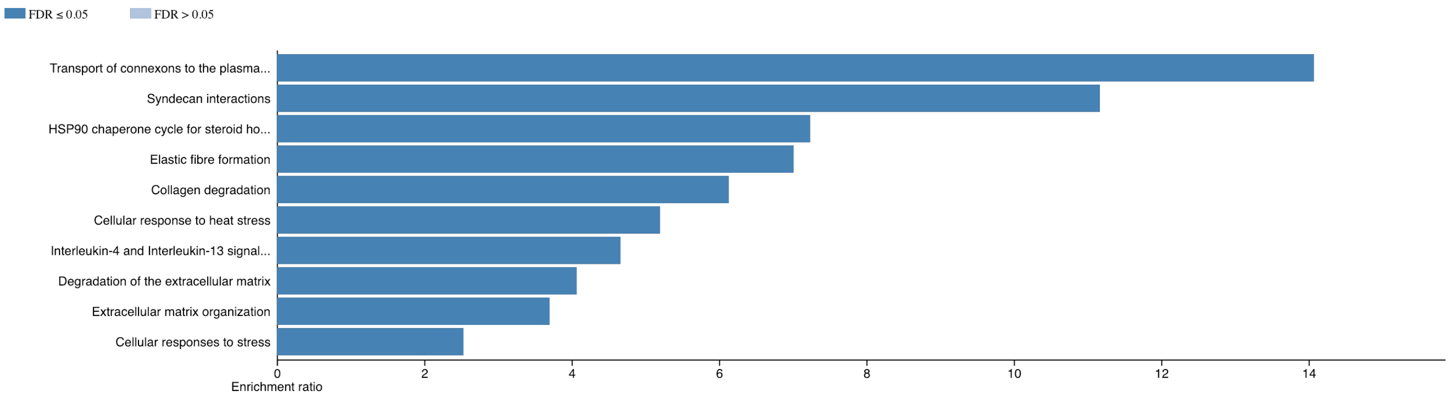

**Figure S14.** Top pathways enriched from 444 genes significantly upregulated in *HAS1*$^{hi}$ fibroblasts compared to other mesenchymal cells.

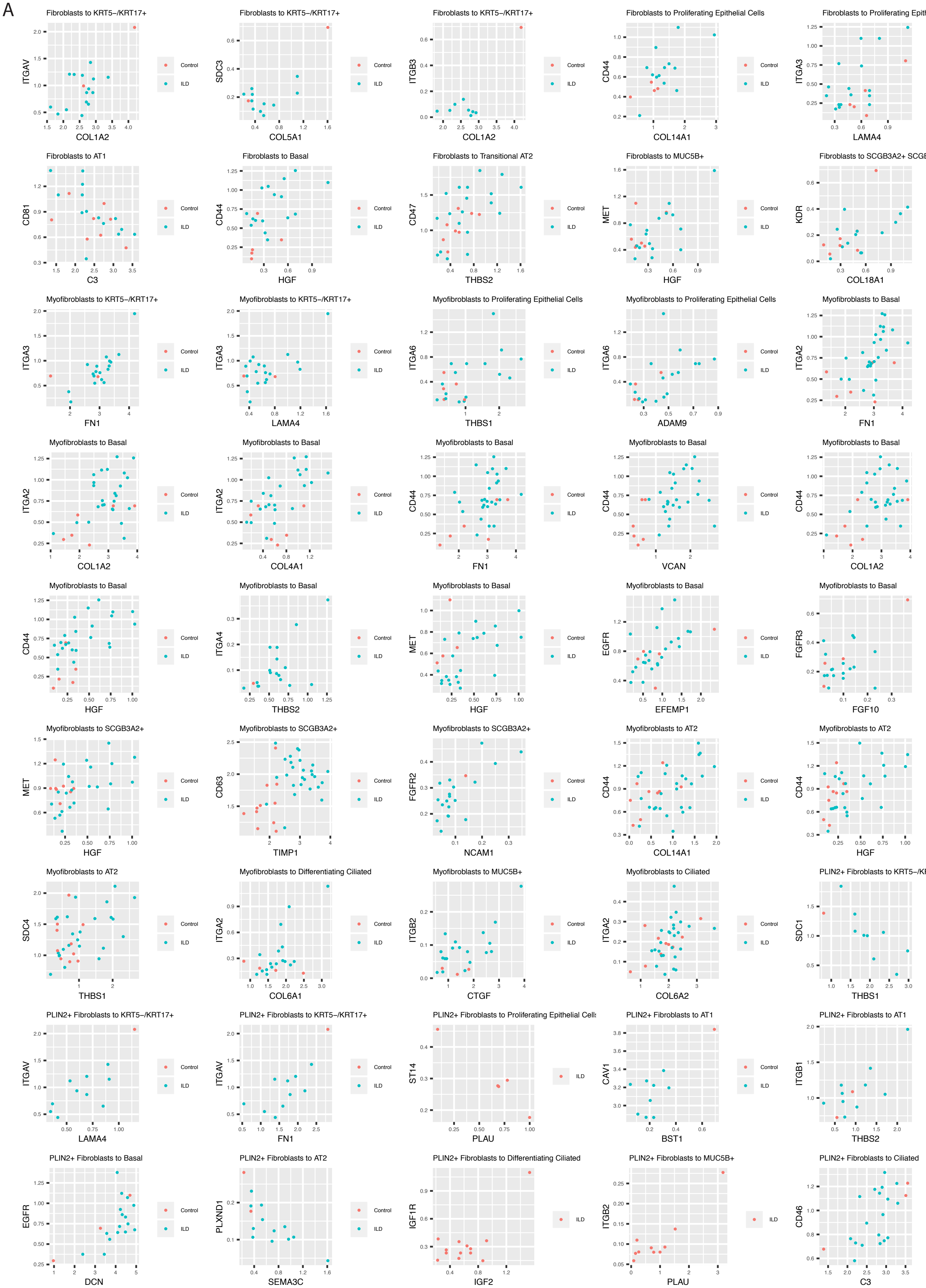
A
Fibroblasts to KRT5−/KRT17+
ITGAV
COL1A2
Fibroblasts to KRT5−/KRT17+
SDC3
COL5A1
Fibroblasts to KRT5−/KRT17+
ITGB3
COL1A2
Fibroblasts to Proliferating Epithelial Cells
CD44
COL14A1
Fibroblasts to Proliferating Epithelial Cells
ITGA3
LAMA4
Fibroblasts to AT1
CD81
C3
Fibroblasts to Basal
CD44
HGF
Fibroblasts to Transitional AT2
CD47
THBS2
Fibroblasts to MUC5B+
MET
HGF
Fibroblasts to SCGB3A2+ SCGB1A1+
KDR
COL18A1
Myofibroblasts to KRT5−/KRT17+
ITGA3
FN1
Myofibroblasts to KRT5−/KRT17+
ITGA3
LAMA4
Myofibroblasts to Proliferating Epithelial Cells
ITGA6
THBS1
Myofibroblasts to Proliferating Epithelial Cells
ITGA6
ADAM9
Myofibroblasts to Basal
ITGA2
FN1
Myofibroblasts to Basal
ITGA2
COL1A2
Myofibroblasts to Basal
ITGA2
COL4A1
Myofibroblasts to Basal
CD44
FN1
Myofibroblasts to Basal
CD44
VCAN
Myofibroblasts to Basal
CD44
COL1A2
Myofibroblasts to Basal
CD44
HGF
Myofibroblasts to Basal
ITGA4
THBS2
Myofibroblasts to Basal
MET
HGF
Myofibroblasts to Basal
EGFR
EFEMP1
Myofibroblasts to Basal
FGFR3
FGF10
Myofibroblasts to SCGB3A2+
MET
HGF
Myofibroblasts to SCGB3A2+
CD63
TIMP1
Myofibroblasts to SCGB3A2+
FGFR2
NCAM1
Myofibroblasts to AT2
CD44
COL14A1
Myofibroblasts to AT2
CD44
HGF
Myofibroblasts to AT2
SDC4
THBS1
Myofibroblasts to Differentiating Ciliated
ITGA2
COL6A1
Myofibroblasts to MUC5B+
ITGB2
CTGF
Myofibroblasts to Ciliated
ITGA2
COL6A2
PLIN2+ Fibroblasts to KRT5−/KRT17+
SDC1
THBS1
PLIN2+ Fibroblasts to KRT5−/KRT17+
ITGAV
LAMA4
PLIN2+ Fibroblasts to KRT5−/KRT17+
ITGAV
FN1
PLIN2+ Fibroblasts to Proliferating Epithelial Cells
ST14
PLAU
PLIN2+ Fibroblasts to AT1
CAV1
BST1
PLIN2+ Fibroblasts to AT1
ITGB1
THBS2
PLIN2+ Fibroblasts to Basal
EGFR
DCN
PLIN2+ Fibroblasts to AT2
PLXND1
SEMA3C
PLIN2+ Fibroblasts to Differentiating Ciliated
IGF1R
IGF2
PLIN2+ Fibroblasts to MUC5B+
ITGB2
PLAU
PLIN2+ Fibroblasts to Ciliated
CD46
C3
Control
ILD

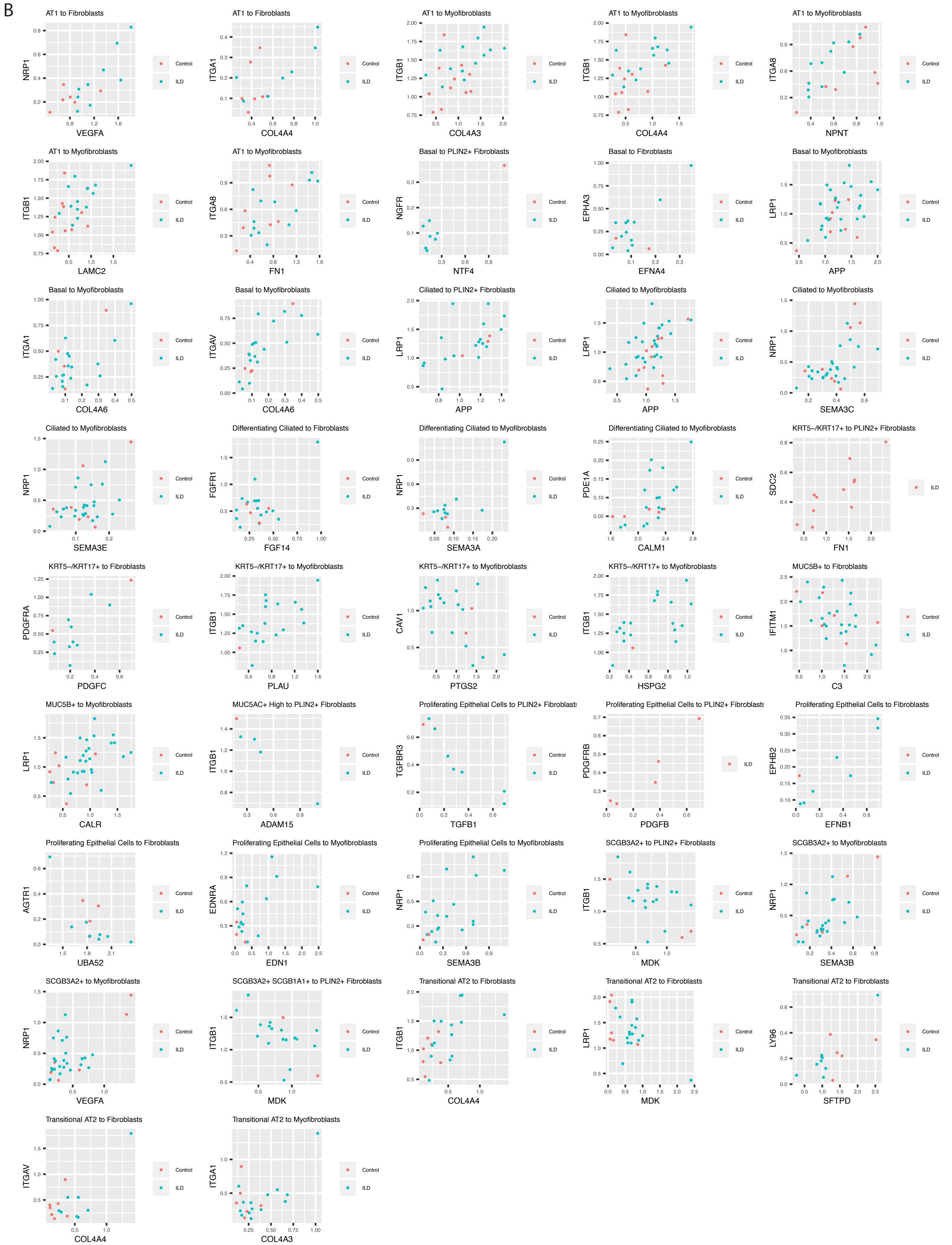

**Figure S15.** Significantly correlated ligand-receptor pair gene expression within individuals for mesenchymal ligands to epithelial receptors (A) and epithelial ligands to mesenchymal receptors (B).

| | Control | PF |
|---|---|---|
| | n=10 | n=20 |
| Age | 37 [17 - 59] | 61.5 [46 -72] |
| Sex (male) | 7 (70%) | 13 (65%) |
| Tobacco | 8 (80%) | 11 (55%) |
| Ancestry | | |
| European | 7 (70%) | 14 (70%) |
| African | 1 (10%) | 4 (20%) |
| Hispanic | | 2 (10%) |
| Other/Unknown | 2 (20%) | 1 (5%) |
| Diagnosis | | |
| IPF | | 12 (60%) |
| NSIP | | 3 (15%) |
| cHP | | 2 (10%) |
| Sarcoidosis | | 2 (10%) |
| Unclassifiable ILD | | 1 (5%) |

**Table S1**. Demographics of lung donors used for scRNA-seq.

| | IPF | Control | NSIP | cHP | Unclassifiable ILD | Sarcoidosis | Total Cells |
|---|---|---|---|---|---|---|---|
| MUC5AC+ High | 75 | 20 | 7 | 1 | - | - | 103 |
| Basal | 1,786 | 65 | 61 | 107 | 37 | 2 | 2,058 |
| Proliferating Epithelial Cells | 222 | 79 | 9 | 20 | 4 | 7 | 341 |
| Differentiating Ciliated | 1,057 | 226 | 8 | 54 | 5 | 12 | 1,362 |
| Ciliated | 11,857 | 2,119 | 169 | 424 | 26 | 76 | 14,671 |
| SCGB3A2+ SCGB1A1+ | 854 | 222 | 58 | 34 | 27 | 215 | 1,410 |
| SCGB3A2+ | 2,824 | 97 | 20 | 145 | 108 | 26 | 3,220 |
| MUC5B+ | 1,940 | 281 | 42 | 112 | 55 | 3 | 2,433 |
| KRT5-/KRT17+ | 374 | 13 | 1 | 62 | 26 | 9 | 485 |
| Transitional AT2 | 581 | 443 | 21 | 84 | 22 | 9 | 1,160 |
| AT2 | 3,225 | 4,393 | 299 | 382 | 498 | 514 | 9,311 |
| AT1 | 96 | 472 | 158 | 15 | 3 | 27 | 771 |
| Lymphatic Endothelial Cells | 584 | 395 | 80 | 68 | 4 | 72 | 1,203 |
| Endothelial Cells | 3,496 | 2,761 | 1,205 | 694 | 290 | 797 | 9,243 |
| Proliferating Macrophages | 813 | 200 | 188 | 293 | 53 | 16 | 1,563 |
| Proliferating T Cells | 132 | 301 | 11 | 22 | 2 | 22 | 490 |
| NK Cells | 957 | 712 | 383 | 74 | 7 | 189 | 2,322 |
| pDCs | 109 | 17 | 27 | 19 | 3 | 13 | 188 |
| cDCs | 1,061 | 151 | 373 | 224 | 154 | 169 | 2,132 |
| Mast Cells | 360 | 254 | 10 | 6 | 36 | 35 | 701 |
| Plasma Cells | 265 | 238 | 45 | 45 | 2 | 9 | 604 |
| B Cells | 550 | 140 | 284 | 13 | 4 | 6 | 997 |
| Monocytes | 2,113 | 2,854 | 354 | 257 | 674 | 513 | 6,765 |
| T Cells | 2,829 | 2,622 | 726 | 304 | 46 | 176 | 6,703 |
| Macrophages | 16,507 | 12,004 | 3,299 | 3,492 | 1,827 | 1,799 | 38,928 |
| PLIN2+ Fibroblasts | 1,167 | 6 | 98 | 6 | 3 | 7 | 1,287 |
| Fibroblasts | 152 | 140 | 19 | 102 | 3 | 11 | 427 |
| HAS1 High Fibroblasts | 179 | - | - | - | - | - | 179 |
| Myofibroblasts | 813 | 196 | 237 | 367 | 45 | 54 | 1,712 |
| Mesothelial Cells | 137 | 35 | 11 | 7 | 15 | 28 | 233 |
| Smooth Muscle Cells | 567 | 188 | 435 | 102 | 27 | 75 | 1,394 |
| Total Cells | 57,682 | 31,644 | 8,638 | 7,535 | 4,006 | 4,891 | 114,396 |

**Table S2**. Distribution of cell types (as a proportion of all cells) in PF compared to control lungs.

| Cell Type | Positive Gene Markers | Negative Gene Markers |
|---|---|---|
| **Epithelial Cells** | EPCAM | |
| AT1 | AGER, PDPN | |
| AT2 | SFTPC, ABCA3, SFTPCD | |
| Transitional AT2 | SFTPC (low), AGER (low) | |
| Basal | KRT5, KRT17 | COL1A1 |
| KRT5-/KRT17+ | KRT17, COL1A1 | KRT5 |
| MUC5B+ | MUC5B, SCGB1A1 | |
| MUC5AC+ High | MUC5AC | |
| SCGB3A2+ | SCGB3A2 | SCGB1A1 |
| SCGB3A2+ SCGB1A1+ | MGP, SCGB1A1, SCGB3A2 | |
| Ciliated | FOXJ1, TMEM190, CAPS, HYDIN | |
| Differentiating Ciliated | FOXJ1, SFTPB | |
| Proliferating Epithelial Cells | MKI67, CDK1 | |
| **Immune Cells** | PTPRC | |
| T Cells | CD3E, FOXP3, IL7R, CD8A, CCL5 | |
| NK Cells | NCR1, KLRB1, NKG7 (high) | CD3E |
| Macrophages | LYZ, MARCO, FCGR1A, C1QA, APOC1 | |
| Monocytes | S100A12, FCN1, S100A9, LYZ, CD14 | |
| cDCs | FCER1A, CD1C, CLEC9A | |
| pDCs | LILRA4, CLEC4C, JCHAIN | |
| Plasma Cells | JCHAIN, IGHG1, IGLL5 | |
| B Cells | MS4A1, CD19, CD79A | |
| Mast Cells | CPA3, KIT | |
| Proliferating T Cells | MKI67, CDK1, CD3E | |
| Proliferating Macrophages | MKI67, CDK1, LYZ | |
| **Endothelial Cells** | PECAM1 | PTPRC |
| Endothelial Cells | VWF, PECAM1 | |
| Lymphatic Endothelial Cells | CCL21 | |
| **Mesenchymal Cells** | | EPCAM, PTPRC, PECAM1 |
| Smooth Muscle Cells | ACTA2 (high), MYH11, PDGFRB (high) | LUM, PDGFRA |
| Mesothelial Cells | WT1, UPK3B | LUM |
| Myofibroblasts | LUM, PDGFRA, ACTA2, MYLK | |
| HAS1 High Fibroblasts | LUM, PDGFRA, HAS1, TWIST | PLIN2 |
| Fibroblasts | LUM, PDGFRA | |
| PLIN2+ Fibroblasts | LUM, PDGFRA, PLIN2 | |

**Table S3**. Marker genes used for cell-type annotation.

**Table S4**. Differentially expressed genes.

**Mesenchymal**: Differential expression analysis between each cell type against all cell types in Mesenchymal population (IPF samples only) and between each diagnosis against control for individual cell types in Mesenchymal population. **KRT5/KRT17_DEG:** Differential expression analysis between KRT5-/KRT17+ cell type against AT1, AT2, Basal, Trans_AT2, Fibroblast, HAS1 High, Myofibroblast and PLIN2+ (Figure S3). **IPF_vs_NonIPF**: Differential expression analysis between IPF and non IPF samples in all cell types (with > 50 cells). **Disease_vs_Control**: Differential expression analysis between disease and control for all cell types (with > 50 cells ; Figure 1). **Control_only_CT_vs_all**: Differential expression analysis between each cell type against all cell types in control samples only. **IPF_only**: Differential expression analysis between HAS1 and KRT5-/KRT17+ cell type vs all mesenchymal cells, fibroblast, mesothelial cells, myofibroblast and PLIN2+ for only IPF samples. **CT_vs_all_per_group**: Differential expression analysis between each cell type against cell type sub groups in in control samples only.

**Table S5.** Differentially expressed genes in KRT5-/KRT17+ cells compared to all epithelial cells, basal cells, AT1 and AT2 cells.

**Table S6.** Variable genes across KRT5-/KRT17+ pseudotime trajectories.

**Table S7**. SOX and NR1D1 motif distribution on promoter of genes in bin III and IV from the AT2_transition_to KRT5-/KRT17+ heatmap.

**See zip file**

**Table S8.** The top 20% highly co-expressed ligand-receptor pairs for each epithelial-mesenchymal interaction from the top 20% highly expressed genes for each cell type (related to Fig. 4 and Fig. S15).
